# Antipsychotic-induced epigenomic reorganization in frontal cortex of individuals with schizophrenia

**DOI:** 10.1101/2021.07.14.452426

**Authors:** Bohan Zhu, Richard I. Ainsworth, Zengmiao Wang, Zhengzhi Liu, Salvador Sierra, Chengyu Deng, Luis F. Callado, J. Javier Meana, Wei Wang, Chang Lu, Javier González-Maeso

## Abstract

Genome-wide association studies have revealed >270 loci associated with schizophrenia risk, yet these genetic factors do not seem to be sufficient to fully explain the molecular determinants behind this psychiatric condition. Epigenetic marks such as post-translational histone modifications remain largely plastic during development and adulthood, allowing a dynamic impact of environmental factors, including antipsychotic medications, on access to genes and regulatory elements. However, few studies so far have profiled cell-specific genome-wide histone modifications in postmortem brain samples from schizophrenia subjects, or the effect of antipsychotic treatment on such epigenetic marks. Here we conducted ChIP-seq analyses focusing on histone marks indicative of active enhancers (H3K27ac) and active promoters (H3K4me3), alongside RNA-seq, using frontal cortex samples from antipsychotic-free (AF) and antipsychotic-treated (AT) individuals with schizophrenia, as well as individually matched controls (n = 58). Schizophrenia subjects exhibited thousands of neuronal and non-neuronal epigenetic differences at regions that included several susceptibility genetic loci, such as *NRG1*, *DISC1,* and *DRD3*. By analyzing the AF and AT cohorts separately, we identified schizophrenia-associated alterations in specific transcription factors, their regulatees, and epigenomic and transcriptomic features that were reversed by antipsychotic treatment; as well as those that represented a consequence of antipsychotic medication rather than a hallmark of schizophrenia in postmortem human brain samples. Notably, we also found that the effect of age on epigenomic landscapes was more pronounced in frontal cortex of AT-schizophrenics, as compared to AF-schizophrenics and controls. Together, these data provide important evidence of epigenetic alterations in the frontal cortex of individuals with schizophrenia, and remark for the first time the impact of age and antipsychotic treatment on chromatin organization.

## Introduction

Schizophrenia has traditionally been viewed as a genetic disorder, with heritability rates estimated at ∼73% (Sawa and Snyder, 2002; Freedman, 2003). However, previous genome-wide association studies (GWAS) clearly showed a relatively low number of genetic regions associated with schizophrenia risk – these include 108 loci in the first study (Schizophrenia Working Group of the Psychiatric Genomics, 2014) that have been expanded to over 270 regions (Farrell et al., 2015; Trubetskoy et al., 2022). Most of these genetic variants were located in non-coding regions and hence, with a few exceptions there is little evidence supporting that coding variants contribute to schizophrenia risk, which also suggests that genetic factors do not seem to be sufficient to fully explain the molecular causes underlying this severe psychiatric condition.

Twin studies have provided evidence that environmental factors contribute to schizophrenia susceptibility. Thus, it had originally been suggested that monozygotic twins, whose DNA sequences are approximately 100% identical, have a concordance for schizophrenia of nearly 50% (Cardno and Gottesman, 2000; Hallmayer et al., 2011), yet recent research has revealed a probandwise concordance rate of 33% in monozygotic twins and 7% in dizygotic twins (Hilker et al., 2018). While these findings underscore the substantial influence of genetic factors on the etiology of schizophrenia, they also advocate for a significant involvement of environmental events in the intricate development of this complex disorder (Li et al., 2021). This concept is further supported by epidemiological studies suggesting that prenatal environmental insults, such as maternal infection (Brown et al., 2004; Yudofsky, 2009) and severe adverse life events (Malaspina et al., 2008), increase the risk of schizophrenia in the offspring.

Gene expression is regulated by the ability of the transcriptional machinery to access DNA, which is tightly packed into chromatin. The status of chromatin organization depends on epigenetic factors, such as DNA methylation and histone modifications that primarily occur on amino-terminal tails (Graff and Tsai, 2013; Onuchic et al., 2018; Bastle and Maze, 2019). Hence these epigenetic mechanisms lead to stable changes in gene expression that are mediated via altered chromatin structure without modification of DNA sequence, and remain largely plastic throughout all periods of brain development and aging. It is then tempting to speculate that epigenetic mechanisms mediate, at least in part, the effects of environmental factors on central nervous system (CNS) gene activity, and are therefore potentially involved in the pathophysiology of schizophrenia and other mental illnesses.

Supporting this concept, previous studies reported alterations in chromatin structure and accessibility in tissue samples from schizophrenic subjects and controls. Most of these previous reports, however, were focused on DNA methylation differences in peripheral blood (Aberg et al., 2014) and brain (Jaffe et al., 2016; Mendizabal et al., 2019). Using the assay for transposase-accessible chromatin sequencing (ATAC-seq) in bulk tissue homogenates of postmortem frontal cortex samples, only a few differences in chromatin accessibility were observed between schizophrenia subjects and controls, in contrast to thousands of age-related differential accessible chromatin regions (Bryois et al., 2018). Histone modifications, including histone H3 acetylation of lysine 27 (H3K27ac) and histone H3 trimethylation of lysine 4 (H3K4me3) are critically involved in epigenomic regulations; H3K27ac marks active enhancers (Creyghton et al., 2010), whereas H3K4me3 marks active promoters (Bernstein et al., 2005). Enhancers are highly dynamic cis-regulatory elements with known involvement in neurodevelopmental processes (Won et al., 2019), and the dynamics of promoters are also significantly connected with the genetic risk of certain psychiatric conditions (Dincer et al., 2015). However, very few studies have been conducted about potential cell-type-specific genome-wide variations in covalent histone modifications in postmortem human brain samples of individuals with schizophrenia.

As an example, recent work combined fluorescence-activated cell sorting (FACS) of neuronal and non-neuronal cell nuclei with chromatin immunoprecipitation sequencing (ChIP-seq) assays in two brain regions (prefrontal cortex and anterior cingulate cortex) from postmortem brain samples of subjects without any known neurological or psychiatric disease (Girdhar et al., 2018). Besides the identification of cell and region-specific histone modification landscapes in this cohort of control subjects, these findings also compared their datasets with previous GWAS of individuals with psychiatric conditions, reporting that strong specific enrichments occurred with schizophrenia and weaker associations with depression in both H3K27ac and H3K4me3 peaks. This correlation was almost exclusively observed in neuronal chromatin, but not in non-neuronal cell nuclei. More recent investigation conducting H3K27ac and H3K4me3 ChIP-seq assays in cortical neurons from schizophrenia subjects and controls has identified rare specific epigenetic variations for a set of non-coding RNA genes (Gusev et al., 2019) and chromatin domain alterations (Girdhar et al., 2022) that may contribute to the pathogenesis of schizophrenia. However, these previous epigenomic studies in postmortem human brain samples did not address the potential effect of previous exposure to antipsychotics on the regulation of chromatin state. This is particularly relevant considering that repeated administration of antipsychotic medications leads to epigenetic modifications at selected gene regions in mouse (de la Fuente Revenga et al., 2018; Ma et al., 2018) and postmortem human brain (Kurita et al., 2012; Ibi et al., 2017) samples. Similarly, whether such type of schizophrenia-associated epigenomic changes is observable in non-neuronal frontal cortex nuclei remains unexplored; even though there is evidence that alterations in the glia may contribute to major psychiatric disorders (Liu et al., 2022).

Combining MOWChIP-seq (Cao et al., 2015b; Zhu et al., 2019) and Smart-seq2 (Picelli et al., 2013) for low-input profiling of H3K27ac and H3K4me3 histone modifications and transcriptomes, respectively, here we present the first dataset with cell type-specific epigenomic and transcriptomic landscapes in postmortem frontal cortex samples from two cohorts of schizophrenics either previously treated or not with antipsychotic medications and control subjects individually matched by sex and age. Importantly, our analyses allow to identify transcription factors (TFs), their regulatees, and genes that may be involved in either the therapeutic effects of antipsychotics or the cause of undesired antipsychotic-induced epigenomic aberrations.

## Results

### Quality assessment for ChIP-seq and RNA-seq datasets in frontal cortex from postmortem human brain samples

Frontal cortex is a brain region involved in processes affected in schizophrenia patients, such as perception, cognition and sensorimotor gating (Andreasen et al., 1994). We selected bilateral frontal cortex (Brodmann area 9) gray matter from 58 brain samples (29 schizophrenia and 29 controls). Control subjects were individually matched based on sex and age, and to a lower degree on postmortem delay (or PMD – time between death and tissue sample collection) (Tables S1 and S2). Nuclei were FACS-sorted using an anti-NeuN antibody as a marker of neuronal nuclei, and NeuN-positive (NeuN^+^) and NeuN-negative (NeuN^-^) nuclei (approximately 60,000 nuclei per NeuN^+^ or NeuN^-^ sample) were collected for ChIP-seq (10,000 nuclei per library) and RNA-seq (6,000 nuclei per library) (Figure 1A). After library preparation and sequencing, our MOWChIP-seq technology generated high-quality ChIP-seq data with average unique reads of ∼11 million and ∼14 million on histone modifications H3K27ac and H3K4me3, respectively (Figure S1A; and Table S3). These yields were comparable to those in our previous studies using mouse frontal cortex and mammalian tissue culture samples (Zhu et al., 2019; de la Fuente Revenga et al., 2021). We generated saturation curves to validate that our sequencing depth is sufficient, and that a further increase in the sequencing depth would not lead to significantly more called peaks (Figure S2). MOWChIP-seq datasets have very low background noise (de la Fuente Revenga et al., 2021) with the fraction of reads in called peaks (FRiP) averagely at 17.35% and 27.59% for our H3K27ac and H3K4me3 profiling, respectively (Figure S1B; Table S3). The PCR bottleneck coefficient (PBC) was calculated to measure library complexity (0.90 and 0.92 for H3K27ac and H3K4me3, respectively), which indicates that most of our ChIP-seq datasets have no or mild bottlenecking (Figure S1C; Table S3).

**Figure 1.**
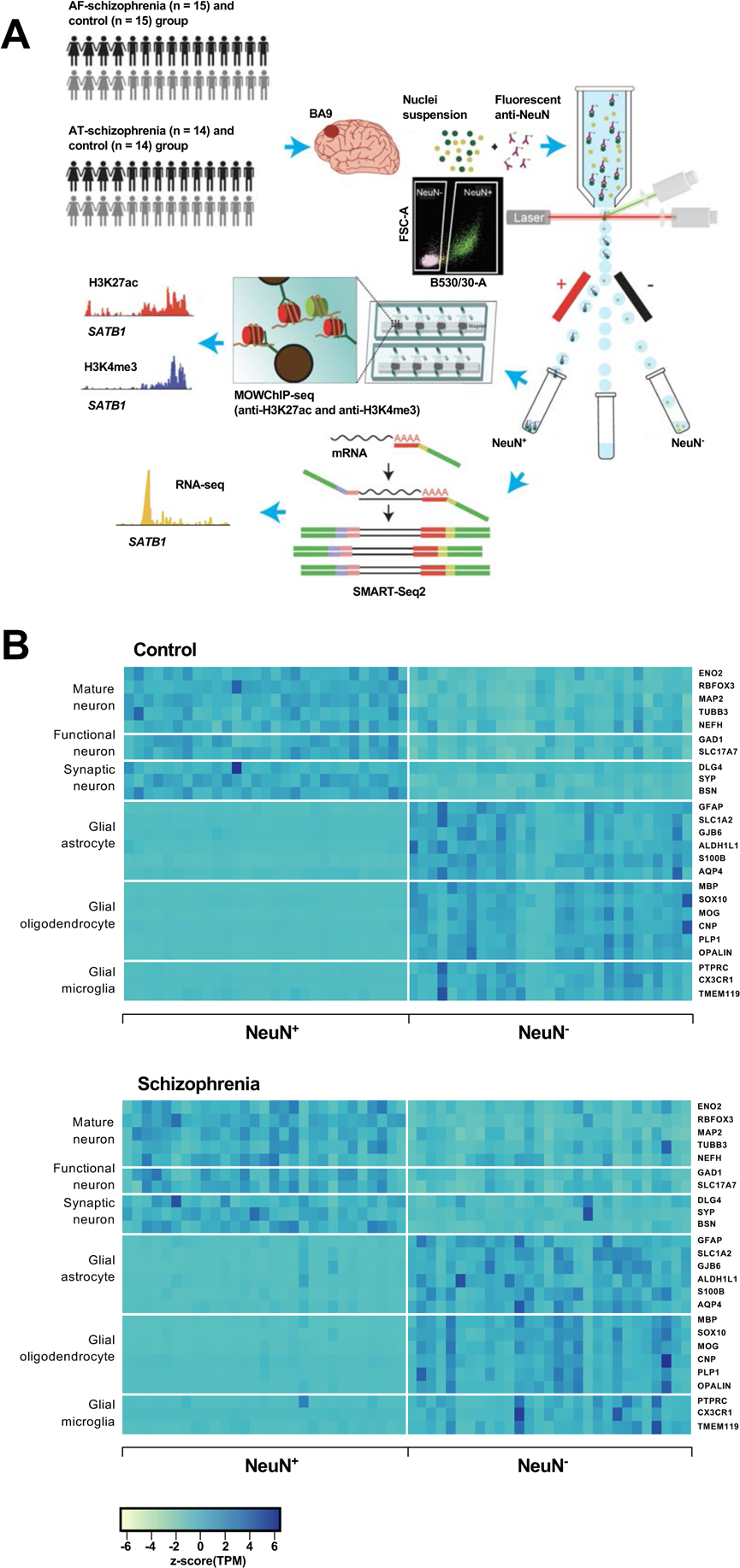
Overview of the multi-omics protocol for analyzing frontal cortex of schizophrenia subjects and controls. (A) Overview of the experimental design starting from postmortem human frontal cortex samples to generate cell type-specific H3K27ac, H3K4me3 and RNA profiles. (B) Heatmap of the expression of neuronal and glial cell markers across all NeuN^+^ and NeuN^-^ frontal cortex samples from 29 control subjects and 29 schizophrenia subjects.

Using *Phantompeakqualtools* (Marinov et al., 2014), we calculated the normalized strand cross-correlation (NSC) and relative strand cross-correlation (RSC) to demonstrate the enrichment of sequencing reads around the histone modification sites (Figures S1D and S1E; Table S3). The average NSC was 1.15 and 1.23 for H3K27ac and H3K4me3, respectively; and the average RSC was 3.75 and 2.09 for H3K27ac and H3K4me3, respectively. These NSC and RSC values were higher than the recommended thresholds of 1.05 and 1.0, respectively. We also compared our GC metrics with those of previously published ChIP-seq data in postmortem human frontal cortex samples (Girdhar et al., 2018). This previous study had an average NSC of 1.33 and 2.81, and RSC of 1.18 and 1.06 for H3K27ac and H3K4me3, respectively. The average Pearson’s correlations between replicates were 0.923 and 0.971 for H3K27ac and H3K4me3, respectively, which compare well with those of ENCODE data (The ENCODE Project Consortium, 2012). Our RNA-seq datasets had an average of ∼ 6.8 million uniquely mapped reads, and the average mapping rate was 82.5% (Figure S3; Table S4), within the recommended range of 70-90% (Conesa et al., 2016). Our average GC content was 40.73% and exon percentage was 36.52%.

Using all frontal cortex samples from the 29 schizophrenia subjects and 29 controls, we analyzed the expression of selected neuronal and non-neuronal marker genes. Highly significant (median p-value = 6 × 10^-7^) pair-wise differences in molecular marker expression were observed for all markers ranging from mature, functional and synaptic neuron markers to astrocyte, oligodendrocyte and microglial markers (Figure 1B; Figures S4 and S5; Table S5) – confirming neuronal and non-neuronal cell-type identities in the NeuN^+^ and NeuN^-^ nuclei samples, respectively.

Consequently, it can be concluded that MOWChIP-seq technology offers data quality comparable to that of state-of-the-art standard reference epigenomes while allowing histone modification profiling using small and highly purified populations of neuronal and non-neuronal nuclei from postmortem human frontal cortex samples.

### Epigenome and transcriptome profiling in neuronal and non-neuronal nuclei from frontal cortex of schizophrenia subjects and controls

To gain insight into the cell type-specific epigenomic changes associated with schizophrenia, we profiled histone marks H3K27ac and H3K4me3 and transcriptomes in NeuN^+^ and NeuN^-^ nuclei from the frontal cortex of schizophrenia subjects and controls. MOWChIP-seq data were mapped to the reference genome (Grch38), and significant peaks were called using MACS2 (Figure S1F). We then performed the overlap analysis to identify consensus peak sets for H3K27ac and H3K4me3 in NeuN^+^ and NeuN^-^ nuclei (see *Methods* section). We identified 107938 consensus peaks covering ∼135 Mb (4.22% of the genome) for H3K27ac in NeuN^+^ nuclei, 71490 consensus peaks covering ∼101 Mb (3.18% of the genome) for H3K27ac in NeuN^-^ nuclei, 133137 consensus peaks covering ∼164 Mb (5.12 % of the genome) for H3K4me3 in NeuN^+^ nuclei, and 100745 consensus peaks covering ∼142 Mb (4.43% of the genome) for H3K4me3 in NeuN^-^ nuclei. This peak distribution was consistent with previous H3K27ac and H3K4me3 NeuN^+^ ChIP-seq studies in postmortem human frontal cortex tissue samples (Girdhar et al., 2022) (Figure S6). Voom plots using raw binding affinity matrix in ChIP-seq and RNA-seq datasets validate that low-enriched/expressed peaks/genes were filtered before downstream analysis (Figure S7). This is further corroborated with the smoothly decreasing curves fitted to the square root of residual standard deviation by average expression in all cases (Figure S7).

We then investigated the differences in histone modification profiles and gene activity between the schizophrenia and control cohorts. We regressed demographic (sex, age at death, PMD, antemortem diagnosis) and technical (align rate, unique rate, FRiP, NSC, RSC, the number of identified peaks and PBC) covariates (Sun et al., 2016). We defined differential H3K27ac peaks that have no overlap regions with promoters as differential enhancers. Our analysis revealed 2301 differential enhancers, 262 differential promoters and 802 differentially expressed genes (DEGs) in NeuN^+^ nuclei between schizophrenia subjects and controls, while 2657 differential enhancers, 360 differential promoters, and 1043 DEGs were discovered in NeuN^-^ nuclei (Figure 2A; Table S6). We next leveraged the previously identified promoter-anchored chromatin loops in NeuN^+^ and NeuN^-^ nuclei to identify how differential enhancers are linked with genes (Hu et al., 2021). We successfully associated 639 and 714 cell-type-specific enhancers to 328 and 395 genes via enhancer-promoter interactions in NeuN^+^ and NeuN^-^ nuclei, respectively. The rest of the enhancers were associated with their nearest genes. We discovered that the schizophrenia group had varied H3K27ac/H3K4me3 levels and RNA-seq reads compared to controls at various loci involved in processes related to synaptic plasticity and cognitive processes or previously associated with schizophrenia risk (Farrell et al., 2015) – these included enhancer region of *NRG1* (*Neuregulin 1*) in NeuN^+^ nuclei, enhancer region of *GRM3* (*Metabotropic glutamate 3 receptor*) in NeuN^-^ nuclei, promoter region of *DRD3 (Dopamine D3 receptor)* in NeuN^+^ nuclei, promoter region of *DISC1* in NeuN^-^ nuclei, *CDK5* (*Cyclin-dependent kinase 5*) mRNA in NeuN^+^ nuclei, and *GRIN2A (Glutamate ionotropic receptor NMDA type subunit 2A)* mRNA in NeuN^-^ nuclei (Figures 2B and 2C; Table S6). The accuracy of H3K27ac and H3K4me3 differential peak calling between schizophrenia subjects and controls identified by MOWChIP-seq was validated by independent ChIP-qPCR analysis of selected loci (Figure S8).

**Figure 2.**
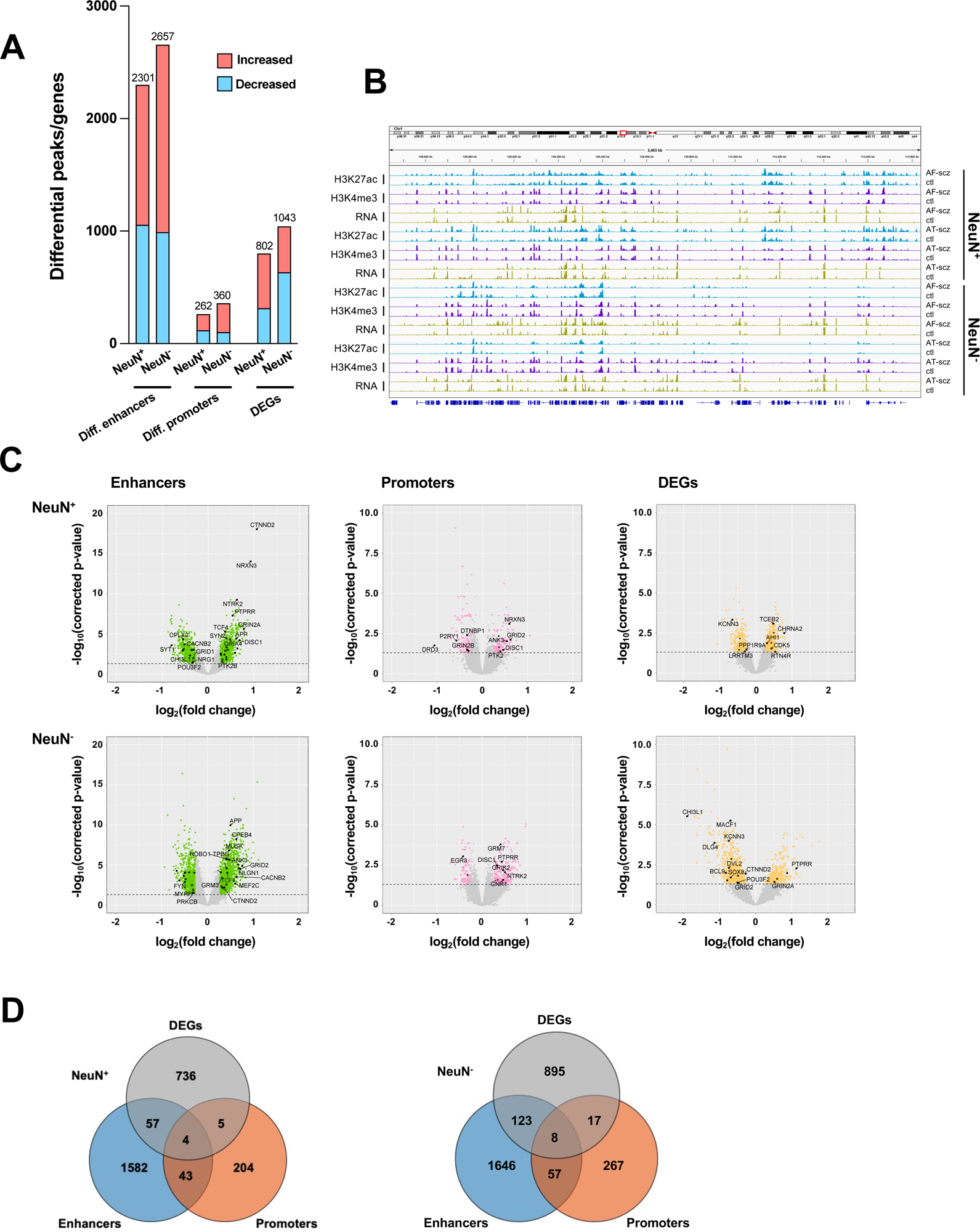
Comparison of epigenomic and transcriptomic landscapes in the frontal cortex of schizophrenia subjects and controls. (A) Differential enhancer/promoter peaks and DEGs obtained by comparing schizophrenia (n = 29) and controls (n = 29). The differential peaks or DEGs were identified using FDR < 0.05. (B) Exemplar genomic track view of H3K27ac, H3K4me3 and RNA signals for matched AF-schizophrenia/control and AT-schizophrenia/control pairs in NeuN^+^ and NeuN^-^ cells. 50Mb region displayed: chr1:68,000,000-118,000,000 (GRCh38). (C) Volcano plots showing genes associated with differential enhancer and promoter peaks and DEGs. Candidate genes for schizophrenia or genes involved in significant GO terms are labeled. The horizontal lines indicate FDR of 0.05. (D) Venn diagrams on the relationship among genes associated with differential enhancer or promoter peaks and DEGs.

We also constructed QQ plots to examine the validity of our differential analysis (Figure S9). The lambda values of our H3K4me3 ChIP-seq datasets were lower than 1 (*i.e*., 0.43 and 0.36 for NeuN^+^ and NeuN^-^ nuclei, respectively), suggesting that there are fewer differential promoters than in the normal distribution. This matches previous findings on postmortem human brain samples (Girdhar et al., 2018).

To further explore the association between differential epigenetic modifications and genetic loci previously associated with schizophrenia risk (Trubetskoy et al., 2022), we examined overlap of differential enhancer and promoter peaks with genetic variants using linkage disequilibrium (LD) score regression (Finucane et al., 2015). Various other brain and non-brain related traits were also considered for comparison (Finucane et al., 2018). Schizophrenia was the most significantly enriched trait for all the differential enhancer and promoter regions in NeuN^+^ and NeuN^-^ fractions (Figure S10). Additionally, the level of enrichment was higher in differential enhancers as compared to promoters, and the highest in differential enhancers of NeuN^+^ nuclei (Figure S10).

To assess agreement with the literature, we compared the DEGs identified in our study with a previous single-nucleus RNA sequencing (snRNA-seq) study in postmortem prefrontal cortex of schizophrenics and controls (Ruzicka et al., 2020). Importantly, 236 out of our 802 DEGs (p-value = 1.96 × 10^-11^) in NeuN^+^ nuclei, and 63 out of our 1043 DEGs (p-value = 4.18 × 10^-6^) in NeuN^-^ nuclei were also identified in this previous single-cell dissection work. In NeuN^+^ nuclei, several genes encoding metabotropic glutamate receptors (*GRM3, GRM5*) that are directly associated with schizophrenia risk (Maj et al., 2016) were found differentially expressed in both studies (Table S7). We also identified some novel genes, including *LRRTM3*, which regulates excitatory synapse development (Um et al., 2016), and *POU3F2*, which is viewed as a key regulator of gene expression in a schizophrenia-associated gene co-expression module (Chen et al., 2018) (Table S7).

We also overlapped genes identified from differential enhancers/promoters with DEGs from RNA-seq (Figure 2D; Table S8). We found that 66 (p-value = 9 × 10^-3^) and 148 (p-value = 1.7 × 10^-2^) genes were identified in both the DEGs from RNA-seq and differential enhancers/promoters associated genes in NeuN^+^ and NeuN^-^ nuclei, respectively. Among these, several schizophrenia-associated genes were also detected, including *NTNG2*, which is known to be involved in neurodevelopmental disorders (Dias et al., 2019) and *GRIN3A*, a gene that encodes NMDA receptor subunits in neuronal nuclei (Yu et al., 2018).

For integrative analysis of these diverse epigenomic and transcriptomic data, we next employed an unbiased method called *EpiSig* (Ai et al., 2018), which combines the epigenomic and transcriptomic dataset into a single analysis to cluster regions with similar epigenomic profiles across all the NeuN^+^ and NeuN^-^ nuclei samples. 85,462 signal enriched regions were grouped into 814 epigenomic clusters covering 14.53% of the genome. These clusters were further combined into 6 groups (sections) using the K-means method (Figures 3A and 3B; Tables S9 and S10). Section I had high coverage in the gene annotations for intron (35%) and intergenic regions (29%) indicating inactive regions. It was also enriched in chromosome X compared to other sections. Section II was annotated as enhancers that are active in NeuN^+^ nuclei but suppressed in NeuN^-^ nuclei. A hypergeometric test identified clusters that were significantly enriched in schizophrenia vs control differential histone marks and differentially expressed genes (Figures 3A and 3B; Tables S9 and S10). The top 5 Section II clusters had schizophrenia positive association (*i.e*., activity and expression schizophrenia > controls) in genes enriched in GO terms “*Trans-synaptic signaling*” (FDR 4.66 × 10^-2^). Section III was highly enriched in enhancers (average of 21% of all regions in each cluster), and low coverage in intergeneric regions (12%), which is likely associated with active enhancers for both NeuN^+^ and NeuN^-^ given the high signal strength for H3K27ac. Using the top differentially enriched cluster genes enriched in the GO term, “*Amyloid fibril formation*” (FDR 5.99 × 10^-2^) was found to be negatively associated with schizophrenia in NeuN^+^ nuclei, whereas genes enriched in the GO term “*Neuron projection development*” (FDR 3.19 × 10^-2^) were positively associated with schizophrenia in NeuN^-^ nuclei (Figures 3A and 3B; Tables S9 and S10). Section IV also had high coverage of enhancers (Figures 3A and 3B; Tables S9 and S10). However, it had the highest average promoter content with 16% of all gene annotations being promoter regions, further supported by CpG islands showing the highest proportion (20%) in this section; indicating active promoters for both NeuN^+^ and NeuN^-^ nuclei. Both cell types showed enrichment in respiratory electron transport genes that were negatively associated with schizophrenia (Figures 3A and 3B; Tables S9 and S10). As for the average differential signals across sections, a great variance was observed. For example, the signal of H3K4me3 was higher in schizophrenia subjects compared to controls for NeuN^+^ nuclei in section V, while it was lower in schizophrenia subjects for NeuN^-^ samples (Figures 3A and 3B; Tables S9 and S10). Finally, Section VI was annotated as enhancers that were active in NeuN^-^ nuclei but repressed in NeuN^+^ nuclei (Figures 3A and 3B; Tables S9 and S10).

**Figure 3.**
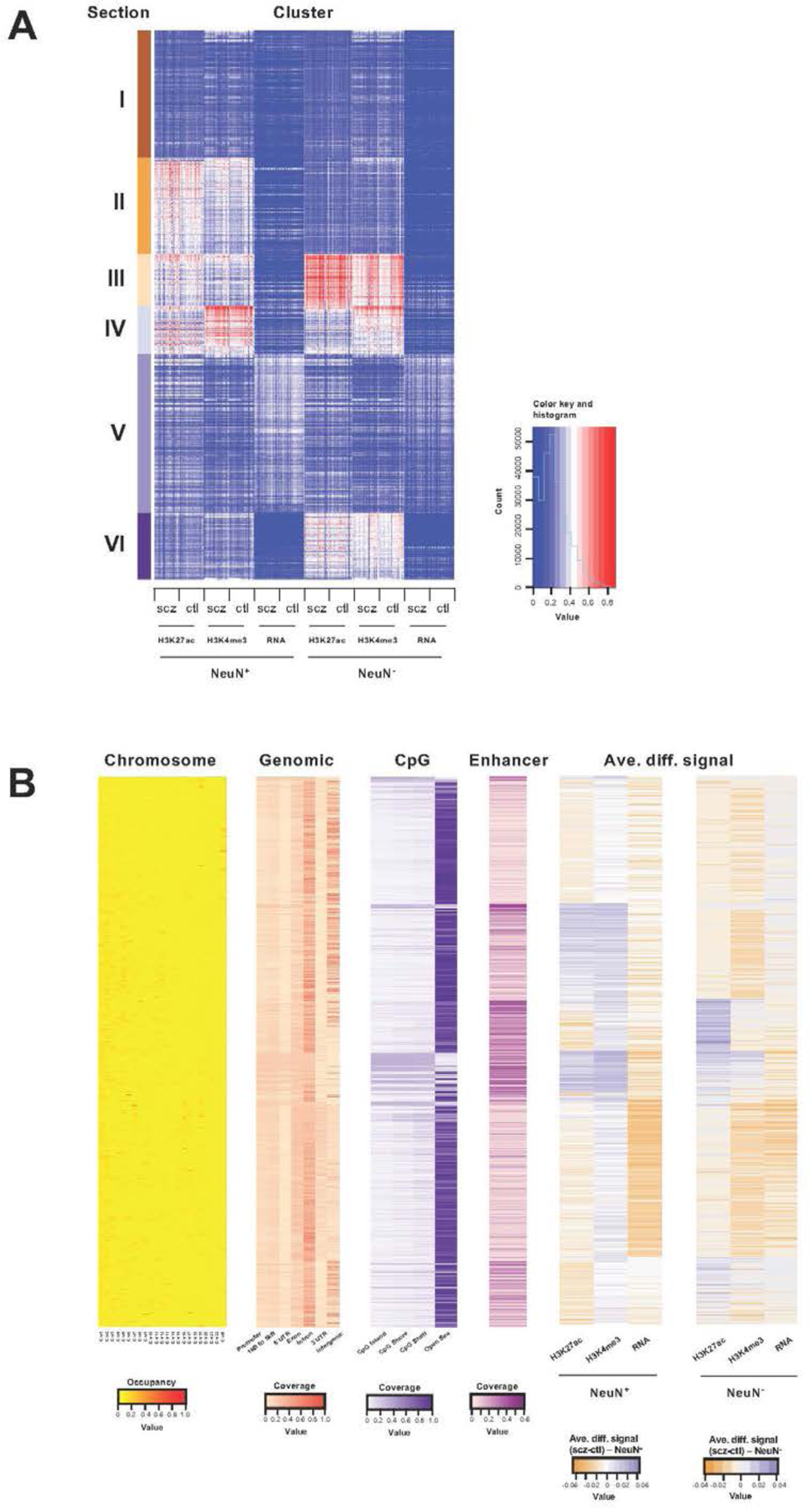
Genome-wide multidimensional clusters in the frontal cortex of schizophrenia subjects and controls. (A) Integrative analysis using *EpiSig*. 814 *EpiSig* clusters across 348 genome-wide sequencing datasets were grouped into 6 sections. The heatmap shows the signal in each *EpiSig* cluster (row: *EpiSig* cluster; column: marker). (B) For each *EpiSig* cluster, from left to right, the heatmaps are: the region percentage in each chromosome; the genomic annotation; the CpG annotation; the percentage of enhancer; the difference signal between schizophrenia and controls in NeuN^+^ and NeuN^-^ nuclei.

Transcriptional regulatory processes proceed as a hierarchy of orchestrated events that ultimately modulate the expression of downstream target genes. Using the recently developed *Taiji* algorithm (Zhang et al., 2019), which allows access to information pertaining to transcriptional cascades deriving from upstream drivers through specific pathway mechanisms to downstream effects, we integrated epigenomic and transcriptomic data to construct 116 individual transcriptional networks in neuronal and glial nuclei from schizophrenia subjects and controls. We identified active promoters and enhancers using H3K27ac and then predicted TF binding sites by scanning 1,165 TF motifs linking putative TF binding sites to their targets using *EpiTensor* (Zhu et al., 2016), an unsupervised method to predict enhancer-promoter associations. TFs were subsequently ranked according to regulatory importance using the Personalized PageRank (PPR) algorithm for each unique network topology (Yu et al., 2017). Using the differentially expressed TFs (schizophrenia *vs* controls FDR < 0.05), TFs were ranked by absolute change in schizophrenia *vs* control PPR score (Figure 4A – top 10 TFs for each cell type; Table S11). Of the top 10 TFs of NeuN^+^ nuclei, all were found to be cell-type specific. Using the top 4 TFs, we identified 207 regulatees the were regulated by 3 or more TFs and found they were involved in processes such as *“Neurexins and neuroligins*” (FDR 2.18 × 10^-7^) and “*Protein-protein interactions at synapses*” (FDR 1.22 × 10^-6^) (Figure 4B; Table S11). Furthermore, all top 10 TFs of NeuN^-^ nuclei were cell-type specific TFs and the regulatees of the top 4 TFs were enriched in signaling pathways including *“RAF/MAP kinase cascade*” (FDR 2.80 × 10^-2^) and “*RHO GTPase cycle*” (FDR 3.07 × 10^-2^) (Figure 4C; Table S11).

**Figure 4.**
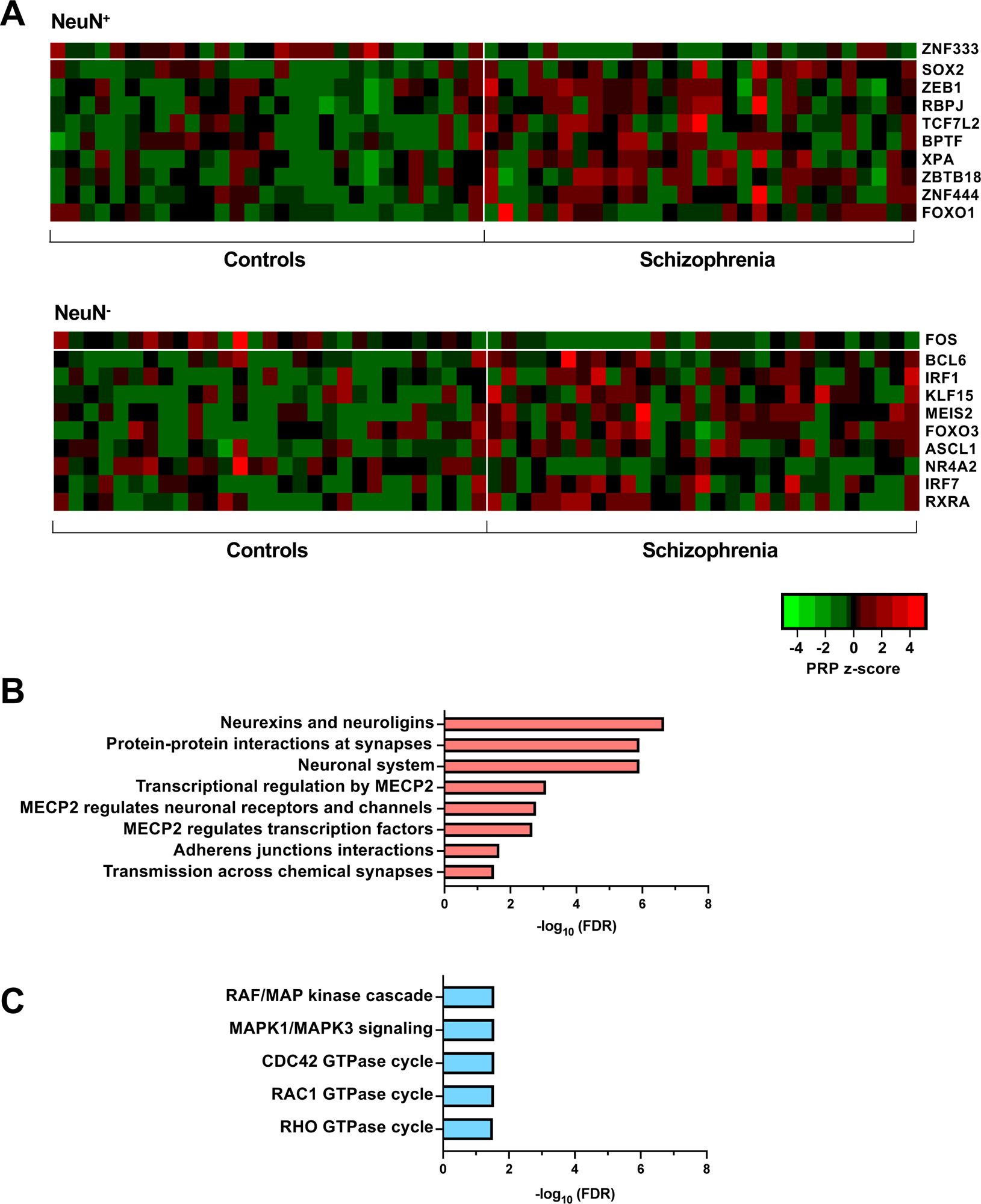
Transcriptional regulatory processes in the frontal cortex of schizophrenia subjects and controls. (A) Heatmap of z-score PPR for top 10 significantly differentially expressed TFs (FDR < 0.05) ranked by absolute change in PPR for NeuN^+^ (upper panel) and NeuN^-^ (lower panel) schizophrenia *vs* control nuclei samples. (B) Overrepresented pathway analysis (FDR < 0.05) for 203 downstream regulatees common to the top 4 schizophrenia *vs* control NeuN^+^-specific TFs (*ZNF333*, *SOX2*, *ZEB1* and *RBPJ*). (C) Overrepresented pathway analysis (FDR < 0.05) for 225 downstream regulatees common to the top 4 schizophrenia *vs* control NeuN^-^-specific TFs (*FOS*, *BCL6*, *IRF1* and *KLF15*).

### Alterations in antipsychotic-free but not in antipsychotic-treated schizophrenics

Using preclinical models, it has been suggested that chronic antipsychotic drug administration leads to long lasting changes in frontal gene expression and chromatin organization (Kurita et al., 2012; de la Fuente Revenga et al., 2018), but the epigenomic consequences of antipsychotic treatment in postmortem human brain samples remain largely unexplored. To validate the separation between the antipsychotic-free (AF) schizophrenia and antipsychotic-treated (AT) schizophrenia groups, we first utilized a dimension reduction algorithm – uniform manifold approximation and projection (UMAP) – to visualize the clustering of each sample with TMM normalized binding affinity matrices or gene expression files from MOWChIP-seq and RNA-seq, respectively (Figure S11). The separation between AF-schizophrenia and AT-schizophrenia groups is clearly visible at enhancer and promoter regions for both NeuN^+^ and NeuN^-^ nuclei.

To further determine the functional relevance of antipsychotic treatment, we aimed to identify the biological pathways, TFs, or gene expressions dysregulated in the AF-schizophrenia group that were also reversed to control levels in the AT-schizophrenia group as compared to individually matched controls. Hence, these represent schizophrenia-associated molecular alterations that are reversed upon antipsychotic treatment. We calculated the average pairwise difference in PageRank in NeuN^+^ nuclei from the AF-schizophrenia/control pair cohort. In conjunction with this, we also calculated the average pairwise difference in PageRank in NeuN^+^ nuclei from the AT-schizophrenia/control pair cohort. We then identified those TFs with a difference in these two values greater than 0.5 (Figure 5A; Table S12). However, when these TFs were further filtered based on a significance cut-off of FDR < 0.05, no significant TFs were identified to be simultaneously changed in the AF-schizophrenia/control group and not changed in the AT-schizophrenia/control group (Figure 5C; Table S12). In parallel, AF-schizophrenia/control DEGs were integrated with genes from the AT-schizophrenia/control cohort with no case/control differences in expression resulting in a list of 116 cohort-specific DEGs (Figure 5C; Table S12). Functional enrichment analysis of these genes resulted in pathways involved in glutamatergic neurotransmission including *“Activation of AMPK downstream of NMDARs”* (FDR 3.66 × 10^-3^) (Figure 5G; Table S12). Structural and functional modifications of dendritic spines are central to brain development and plasticity (Spruston, 2008). Studies from postmortem brains of subjects with neurodevelopmental disorders including schizophrenia demonstrate altered density and morphology of dendritic spines, particularly in the frontal cortex (Glantz and Lewis, 2000; Black et al., 2004). IQGAP scaffold proteins facilitate the formation of complexes that regulate cytoskeletal dynamics including microtubules (Cao et al., 2015a). Interestingly, another significant pathway restored in the AT-schizophrenia group was “*Rho GTPases activate IQGAPs*” (FDR 3.66 × 10^-3^) (Figure 5G; Table S12). The importance of this pathway was validated by the analysis of the clusters from the *EpiSig* pipeline. Thus, taking the top 3 clusters ranked for enrichment in H3K27ac and mapping their differential peaks to genes resulted in 166 genes enriched in pathways including *“Adherens junctions interactions”* (p-value 1.22 × 10^-4^) (Figure S12; Table S13).

**Figure 5.**
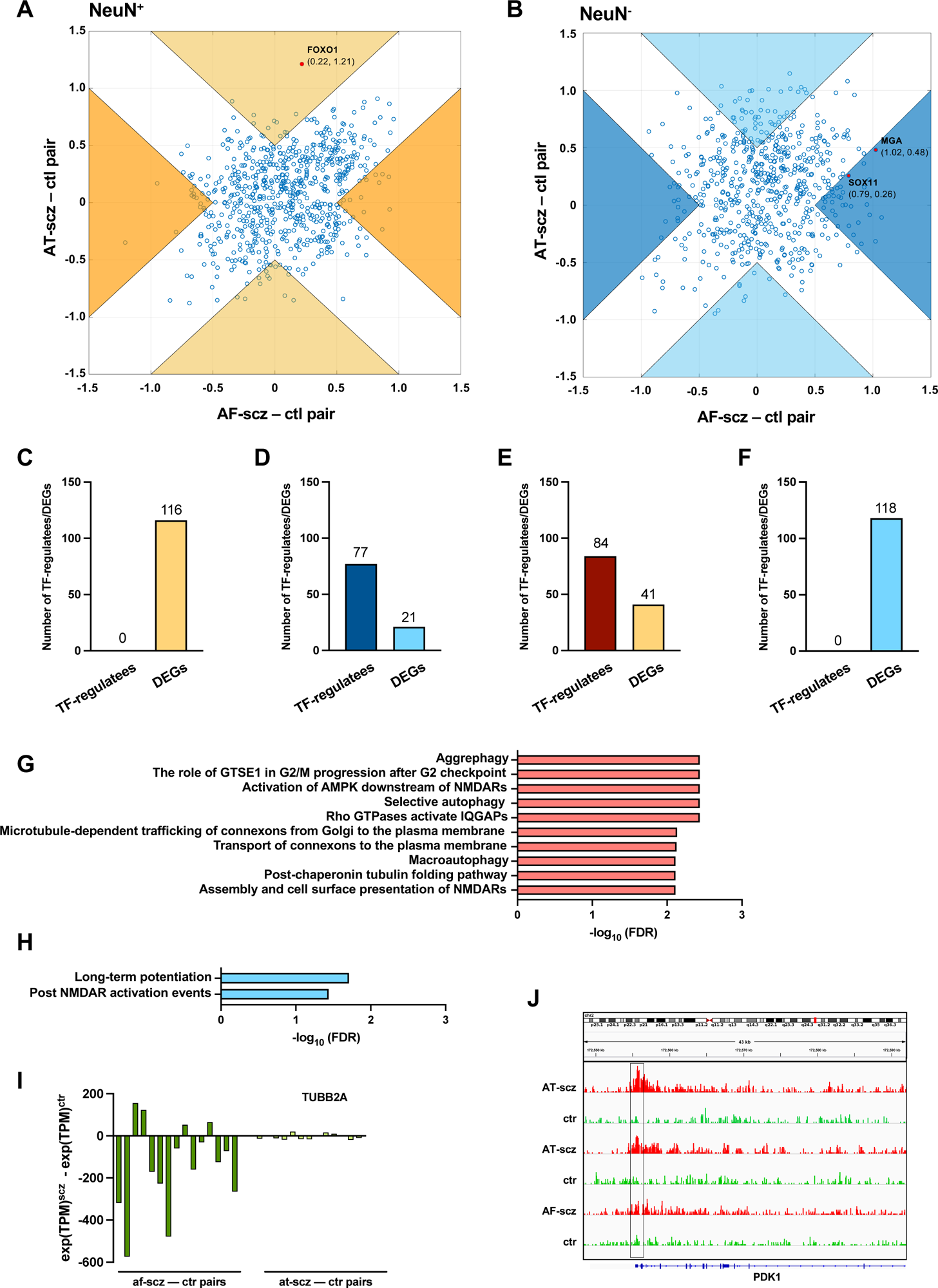
Epigenomic alterations affected by antipsychotic treatment. (A) Scatter plot of average pairwise change in PPR (PPR_schizophrenia_ – PPR_control_) for AF vs AT NeuN^+^ cohorts. Orange regions show cohort (AF – AT) < 0.5 (*i.e*., alterations recovered by antipsychotic treatment), whereas beige regions show cohort (AT – AF) > 0.5 (*i.e*., alterations consequence of antipsychotic treatment). TFs FDR < 0.05 highlighted in red. (B) Scatter plot of average pairwise change in PPR (PPR_schizophrenia_ – PPR_control_) for AF vs AT NeuN^-^ cohorts. Dark blue regions show cohort (AF – AT) < 0.5 (*i.e*., alterations recovered by antipsychotic treatment), whereas cyan regions show cohort (AT – AF) > 0.5 (*i.e*., alterations consequence of antipsychotic treatment). TFs FDR < 0.05 highlighted in red. (C) Number of DEG regulatees by TFs, and number of DEGs in NeuN^+^ nuclei from AF-schizophrenia/control pairs. (D) Number of DEG regulatees by TFs, and number of DEGs in NeuN^-^ nuclei from AF-schizophrenia/control pairs. (E) Number of DEG regulatees by TFs, and number of DEGs in NeuN^+^ nuclei from AT-schizophrenia/control pairs. (F) Number of DEG regulatees by TFs, and number of DEGs in NeuN^-^ nuclei from AT-schizophrenia/control pairs. (G) Functional enrichment analysis of union of genes from AF-schizophrenia/control pairs in NeuN^+^ nuclei. (H) Functional enrichment analysis of union of genes from AF-schizophrenia/control pairs in NeuN^-^ nuclei. (I) Pairwise expression difference (schizophrenia – control) of an exemplar AF-schizophrenia/control cohort DEG (TUBB2A) across all 29 schizophrenia-control pairs in NeuN^+^ nuclei. (J) H3K27ac tracks for PDK1 (member of the 84 gene set in E) in NeuN^+^ nuclei. Box highlighting the FOXO1 DNA-binding motif in promoter at position chr2: 172,555,706 – 172,555,718 (GRCh38). Two exemplar AT-schizophrenia/control cohort pairs showing differential H3K27ac peak intensity around motif locus and an example AF-schizophrenia/control cohort patient pair with no difference.

Autophagy has been suggested to play an important role in the pathophysiology of schizophrenia and antipsychotics are known to modulate the process (Merenlender-Wagner et al., 2015). Notably, the pathways *“Aggrephagy”* (FDR 3.66 × 10^-3^) and *“Macroautophagy”* (FDR 7.71 × 10^-3^) were significantly enriched (Figure 5G; Table S12). Expression of the *“Macroautophagy”* genes *AMBRA1*, *PRKAB1*, *TUBA1A*, *TUBB2A,* and *TUBA4A* was restored in the AT-schizophrenia group (Figure 5I; Table S12). Ubiquitin B (UBB) expression has been previously identified as a strong correlate of schizophrenia symptoms (Rubio et al., 2013). We show a 1.96-fold decrease specific to AF-schizophrenics compared to controls (p-value = 4.0 × 10^-2^) (Table S12).

Glial cells modulate and act as effectors in neurodevelopment through a wide range of neuronal-glial cell interactions. Using the same process as above, we identified 2 driver TFs with significant change in PPR between case and control for the AF-schizophrenia cohort and no significant difference between case and control in the AT-schizophrenia group, *SOX11* (FDR 1.73 × 10^-2^) and *MGA* (FDR 9.83 × 10^-3^) (Figure 5B; Table S12). 77 downstream DEG regulatees of these two TFs were identified in the AF cohort showing significant regulatory case/control change (Figure 5D; Table S12). In parallel, 21 cohort-specific DEGs were identified as AF-schizophrenia/control DEG and having no significant difference in expression in the AT-schizophrenia/control cohort (Figure 5D; Table S12). Functional enrichment analysis of the union of 153 genes included “*Post NMDA receptor activation events* (FDR 3.65 × 10^-2^), and “*Long-term potentiation*” (FDR 1.95 × 10^-2^) (Figure 5H; Table S12). *EpiSig’*s analysis did not show NeuN^-^ alterations in the AF-schizophrenia cohort (data not shown).

We also performed the differential analysis with demographic and technical covariates regressed out on AF-schizophrenia/control and AT-schizophrenia/control cohorts. In NeuN^+^ nuclei, the results revealed 2069 and 574 differential enhancers and promoters, respectively, and 166 DEGs between AF-schizophrenia and their controls (Figure 6A; Table S14), while 3658, 36, and 1273 differential enhancers, promoters, and DEGs were discovered between AT-schizophrenia and controls (Figure 6B; Table S14). In NeuN^-^ nuclei, we identified 891, 19, and 128 differential peaks/genes between AF-schizophrenia and controls (Figure 6A; Table S14); 2651, 775, 776 differential peaks/genes between AT-schizophrenia and controls, in enhancers, promoters, and DEGs, respectively (Figure 6B; Table S14). More differential enhances/promoters and genes were detected between AT-schizophrenia and their matched controls than those between AF-schizophrenia and their controls with the exception in neuronal promoters (Figures 6A and 6B; Table S14).

**Figure 6.**
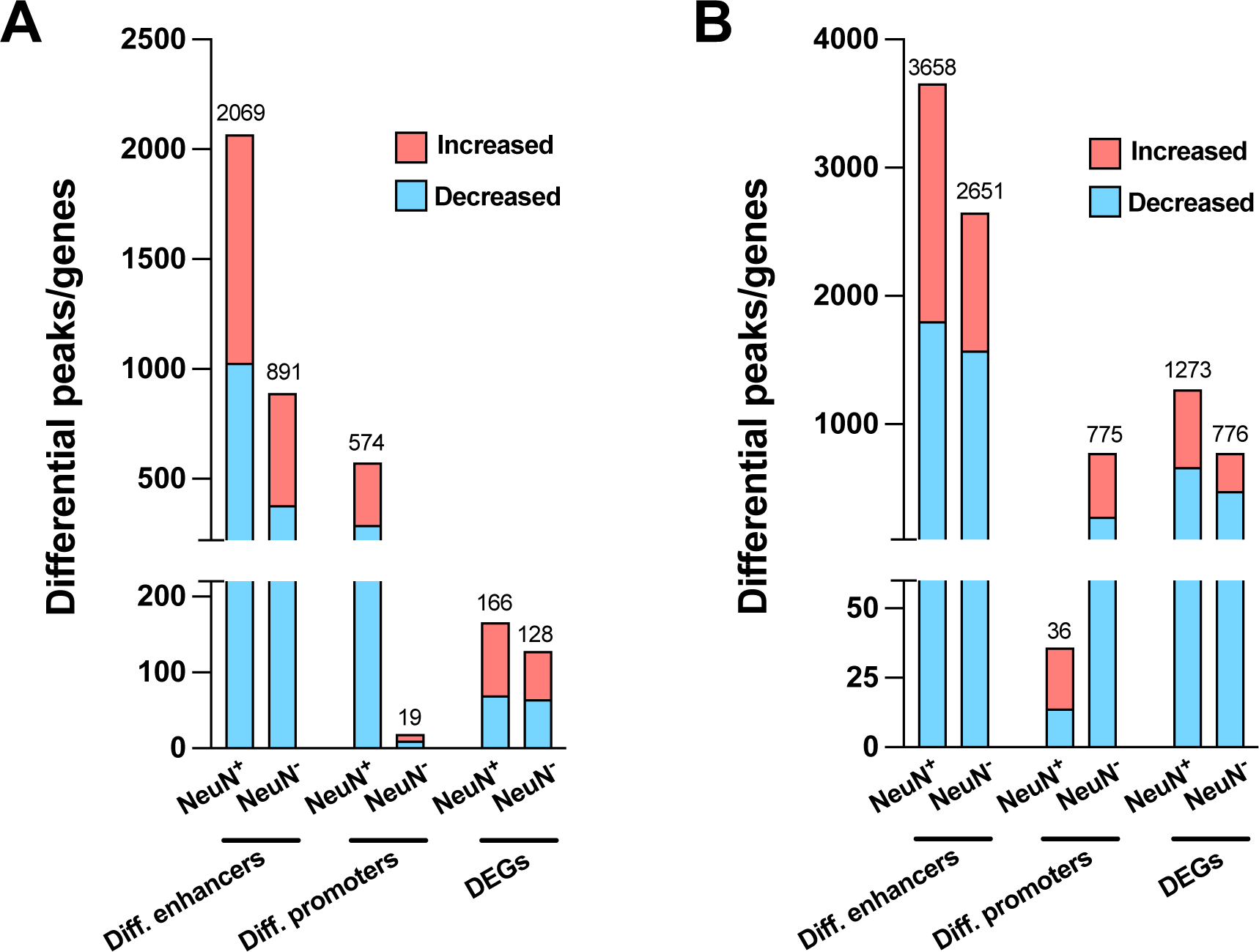
Effect of antipsychotic treatment on differential enhancers/promoters and DEGs in NeuN^+^ and NeuN^-^ nuclei from the frontal cortex of schizophrenia subjects and controls. (A) Differential enhancer/promoter peaks and DEGs obtained by comparing AF-schizophrenics and individually matched controls. (B) Differential enhancer/promoter peaks and DEGs obtained by comparing AT-schizophrenics and individually matched controls.

Similar to our TF analyses (Figure 5), we also identified the genes altered in the AF-schizophrenia/control group but not in the AT-schizophrenia/control group using differential analyses of enhancers, promoters, or gene expression. It should be noted that in the differential analyses here, the schizophrenia subjects (whether AF or AT) and their controls were compared at the cohort level, while matched schizophrenia/control pairs were examined individually in the TF-based analysis. At the epigenomic level, in NeuN^+^ nuclei, we identified 687 and 549 genes changed in the AF-but not in AT-schizophrenics by examining differential enhancers and promoters, respectively (Table S15). These genes were linked to epigenomic features restored to their basal level after treatment. In NeuN^-^ nuclei, there were 270 and 17 recovered genes linked with differential enhancers and promoters, respectively. At the transcriptomic level, 145 DEGs in NeuN^+^ nuclei and 109 in NeuN^-^ nuclei were discovered in AF-schizophrenia/control comparison but not in the AT-schizophrenia/control differential analysis.

### Alterations in antipsychotic-treated but not in antipsychotic-free schizophrenics

We next sorted to characterize those TFs that exhibit regulatory alterations in the AT-schizophrenia/control cohort but not in the AF-schizophrenia/control cohort. Our goal was to identify modifications in pathways that represent a consequence of antipsychotic medication rather that an epigenetic mark of schizophrenia in postmortem human brain (in other words, unwanted side effects caused by antipsychotic treatment). In the same way as above, we identified those TFs with a change in the AT-schizophrenia/control group but not in the AF-schizophrenia/control group for the NeuN^+^ nuclei (Figure 5A; Table S12). Further filtering of these TFs based on a significance cut-off of FDR < 0.05 leads to the identification of FOXO1 (FDR 4.89 × 10^-2^).

We identified dysregulated AT-schizophrenia/control DEG regulatees of these TFs in NeuN^+^ nuclei via analysis of differential edge weights thus obtaining 84 genes (Figure 5E; Table S12). AT-treated/control DEGs were intersected with genes from the AF-schizophrenia/control cohort with no case/control differences in expression resulting in a list of 41 cohort-specific DEGs (Figure 5E; Table S12). Pathway analysis on the union of genes yields the pathway *“Regulation of p53 activity through phosphorylation”* (FDR 1.13 × 10^-2^) (Table S12), including the FOXO1 AT-schizophrenia/control cohort DEG regulatees *CCNA1, BLM, TP53RK*, and *RBBP8*, and the AT-schizophrenia/control cohort-specific DEGs *PRKAA1* and *TAF15* (Table S12). The p53 regulatory gene PDK1 was also identified as a FOXO1 AT-schizophrenia/control DEG regulatee (Figure 5J; Table S12). p53 is one of the most critical pro-apoptotic genes, and antipsychotics are known to produce complex effects including the activation of both proapoptotic and antiapoptotic signaling pathways (Aylon and Oren, 2007). Our data showed that all genes involved in the regulation of p53 presented a significantly lower expression in AT-schizophrenics compared to controls, suggestive of a repressive role for FOXO1 for its 5 regulatees, as FOXO1 had higher PPR in treated schizophrenics (Table S12) and was also 2.38-fold more highly expressed in schizophrenics for the treated-cohort (p-value = 1.07 × 10^-2^) (Table S12). Analysis of the H3K4me3 enriched clusters from the *EpiSig* pipeline for the AT-schizophrenia/control cohort corroborated alterations in pathways related to p53 (Endo et al., 2008) (Figure S13; Table S16).

In NeuN^-^ nuclei, no TFs showed a significant difference in the AT-schizophrenia/control cohort but not in the AF-schizophrenia/control cohort (Figure 5F; Table S12). Furthermore, the 118 genes cohort-specific DEGs identified as AT-schizophrenia/control DEGs and having no significant difference in expression in the AF-schizophrenia/control cohort were not significantly enriched in any signaling pathway. Analysis of the clusters from the *EpiSig* pipeline remarked the importance of the RHO GTPase pathway on the regulatory alterations observed in AT-schizophrenia subjects (Figure S14; Table S17).

We also used differential analyses of enhancers, promoters, and expression to discover the genes altered in the AT-schizophrenia/control group but not in the AF-schizophrenia/control group. In NeuN^+^ nuclei, we found 1591 and 28 treatment-altered genes linked with differential enhancers and promoters, respectively (Table S15). In NeuN^-^ nuclei, we identified 1351 and 718 altered genes linked with differential enhancers and promoters, respectively (Table S15). At the transcriptomic level, 1252 DEGs in NeuN^+^ nuclei and 757 in NeuN^-^ nuclei were discovered in AT-schizophrenia/control comparison but not in the AF-schizophrenia/control differential analysis.

### Age differentially affects antipsychotic-treated schizophrenia subjects

In order to further assess the effect of age on gene expression, we first compared transcriptomes of subjects with schizophrenia and the controls to evaluate how these changes correlated with age. Within NeuN^+^ nuclei in the control group, we identified 742 genes that were significantly correlated with age – with most of them (573, or 77.2%) showing decreased expression in older control subjects (Figure 7A; Table S18). These included *APOL2*, which has been involved in epigenetic aging (Luo et al., 2020). The opposite, however, was observed in NeuN^+^ nuclei from schizophrenia subjects with 18 out of 622 (2.8%) in AF-schizophrenia presenting a negative correlation with age, an effect that was partly reversed in the AT-schizophrenia cohort (85 out of 242 or 35.1%) (Figure 7A; Table S18).

**Figure 7.**
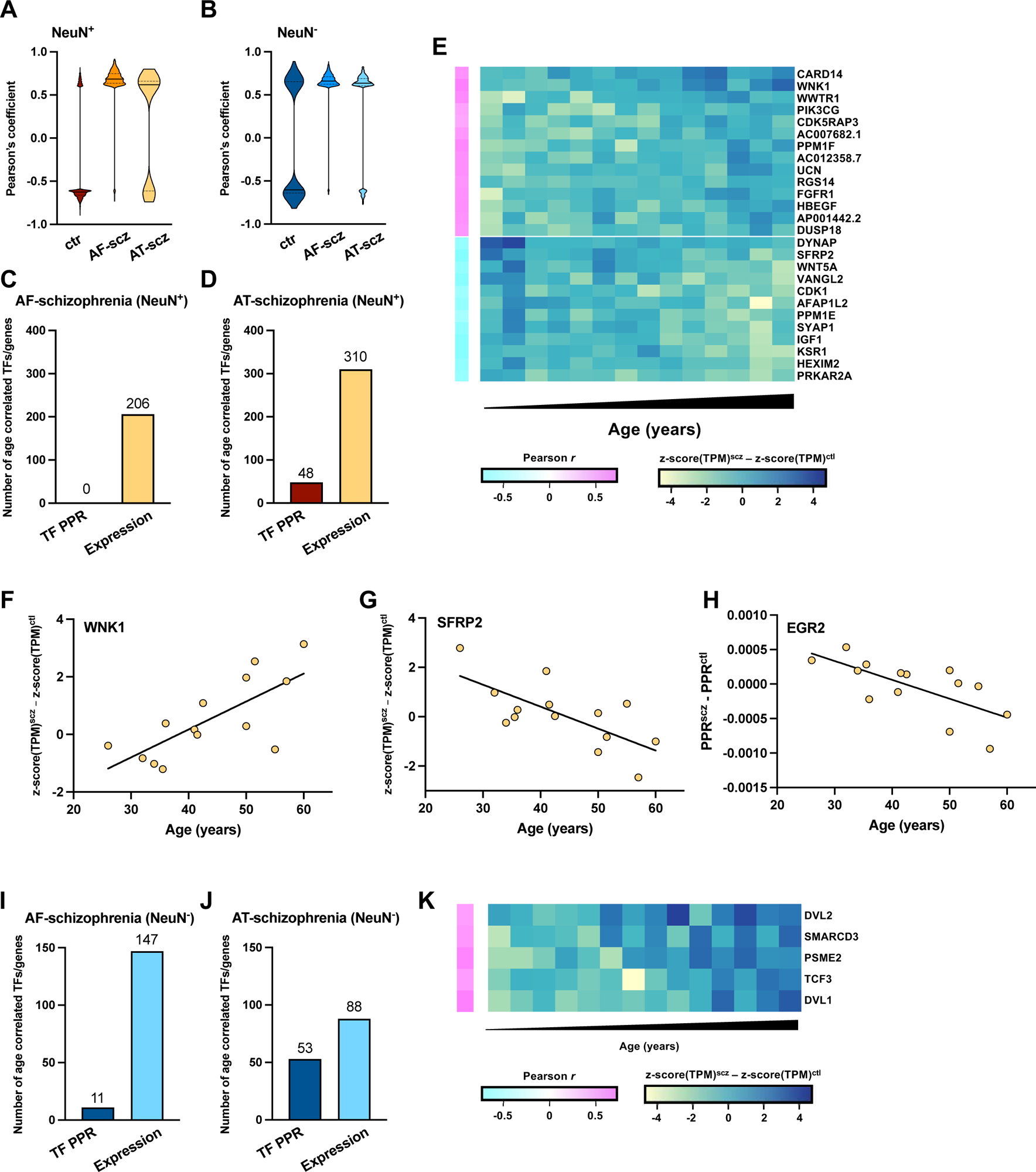
Epigenomic effect of age on treated schizophrenia subjects. (A) Violin plots for Pearson’s R correlation coefficients of age *vs* expression for 742, 622 and 242 genes from control, AF-schizophrenia, and AT-schizophrenia NeuN^+^ nuclei, respectively. (B) Violin plots for Pearson’s R correlation coefficients of age *vs* expression for 1031, 389 and 351 genes from control, AF-schizophrenia, and AT-schizophrenia NeuN^-^ nuclei, respectively. (C) Number of pairwise TF PPR and pairwise gene expression differences correlated with age in NeuN^+^ nuclei from AF-schizophrenia/control pairs. (D) Number of pairwise TF PPR and pairwise gene expression differences correlated with age in NeuN^+^ nuclei from AT-schizophrenia/control pairs. (E) Heatmap for the 14 age positively-correlated (schizophrenia – control increase with age) and 12 age negatively-correlated (schizophrenia – control decrease with age) genes of the significant GO term “*Regulation of kinase activity*” from the AT-schizophrenia/control NeuN^+^ cohort. (F) Example gene, WNK1 pairwise expression difference (schizophrenia – control) vs age (Pearson’s R = 0.73; p-value = 0.003). (G) Example gene, SFRP2 pairwise expression difference (schizophrenia – control) vs age (Pearson’s R = -0.70; p-value = 0.005). (H) Example TF, EGR2 pairwise PPR difference (schizophrenia – control) vs age (Pearson’s R = 0.69; p-value = 0.0003 (I) Number of pairwise TF PPR and pairwise gene expression differences correlated with age in NeuN^-^ nuclei from AF-schizophrenia/control pairs. (J) Number of pairwise TF PPR and pairwise gene expression differences correlated with age in NeuN^-^ nuclei from AT-schizophrenia/control pairs. (K) Heatmap for the 5 age positively-correlated (schizophrenia – control increase with age) genes of the significant GO term “*Beta-catenin independent WNT signaling*” from the AT-schizophrenia/control NeuN^-^ cohort.

Our data also demonstrate that within the NeuN^-^ nuclei genes correlated with age (1031, 389 and 351 in controls, AF-schizophrenia and AT-schizophrenia, respectively), approximately half (491 or 47.6%) were positively correlated with age in the control group whereas a much higher fraction of genes showed increased expression with age in the schizophrenia group, particularly in the AF-schizophrenia cohort (382 or 98.2% in AF-schizophrenia, and 276 or 78.6% in AT-schizophrenia) (Figure 7B; Table S18). These results suggest that age differentially affects gene expression in the frontal cortex of AF-schizophrenia vs AT-schizophrenia subjects as compared to age-matched controls. Importantly, this was further confirmed by functional integration of epigenomic and transcriptomic data and the evaluation of how these alterations correlated with age.

Thus, we evaluated pairwise changes in expression between schizophrenia subjects and their age-matched controls, and identified 206 and 310 genes with an absolute Pearson’s correlation of ≥ 0.50 in NeuN^+^ nuclei from the AF-schizophrenia/control and AT-schizophrenia/control cohorts, respectively (Figures 7C and 7D; Table S18). We also found enriched biological processes associated with age, including “*Regulation of protein kinase activity*” (p-value 6.69 × 10^-7^) in the AT-schizophrenia/control group (Figure 7E; Table S18). Within this gene set, the difference between AT-schizophrenia subjects and control pairs correlated either positively (*WNK1*) or negatively (*SFRP2*) with age (Figures 7F and 7G; Table S18). Evaluation of pairwise changes in PPR identified 48 TFs with high correlations to age in NeuN^+^ nuclei from AT-schizophrenia/control cohorts (Figure 7D; Table S18), whereas this alteration was not observed in the AF-schizophrenia/control group (Figures 7C). Pathway analysis of the NeuN^+^ TFs affected by age in the AT-schizophrenia/control cohort led to the top pathway “*NGF-simulated transcription*” (p-value 8.04 × 10^-8^), including the TFs *EGR2* and *ATF2* (Table S18). Hallucinations and delusions typically attenuate with aging (Davidson et al., 1995), which is consistent with the lower PPR difference for *EGR2* – a preclinical marker of psychosis-like behavior (Gonzalez-Maeso et al., 2007) – that we observed in older subjects (Figure 7H; Table S18).

In NeuN^-^ nuclei, 147 and 88 genes were identified with AF-schizophrenia/control and AT-schizophrenia/control expression difference vs age correlations of ≥ 0.60, respectively (Figures 7I and 7J; Table S18). Enriched pathways in the AT-schizophrenia/control group included: *“Degradation of DVL”* (p-value 4.04 × 10^-5^) and *“Beta-catenin independent WNT signaling”* (p-value 5.06 × 10^-4^) (Figure 7K; Table S18). Since dysfunctional WNT signaling is associated with several CNS disorders including Alzheimer’s (Wan et al., 2014), together, these data also suggest that this positive correlation between NeuN^-^ gene differences in AT-schizophrenia subjects/control pairs and age (Figure 7K; Table S18) may be responsible for some of the negative effects of antipsychotic treatment on cognitive processes. We also identified 11 and 53 TFs correlated with age in the AF-schizophrenia/control and AT-schizophrenia/control cohorts, respectively (Figures 7I and 7J; Table S18). However, as in NeuN^+^ nuclei, the effect of age became more evident in the AT-schizophrenia/control group with age-related adaptations in NeuN^-^ TF-affected pathways that included “*Signaling by NOTCH*” (p-value 3.2 × 10^-4^) (Table S18).

## Discussion

Understanding the molecular determinants involved in schizophrenia is critical for devising new treatment strategies and the discovery of the pathogenic mechanisms underlying this psychiatric condition. In this study, we combined low-input epigenomic and transcriptomic analysis to define how gene expression and TF regulation vary in schizophrenia subjects relative to controls and in response to antipsychotic treatment and aging. Our data provide evidence suggesting alterations in covalent histone modifications at different gene regions previously associated with schizophrenia risk, as well as additional genes involved in pathways related to immunological and neurodevelopmental processes. Whereas previous studies with postmortem human brain samples compared using indirect methods differences in chromatin accessibility between schizophrenia subjects and controls (Bryois et al., 2018), here we provide epigenomic signatures that distinguish between those observed in AF-schizophrenia subjects as well as alterations that denote previous treatment with antipsychotic medications.

We conducted pairwise comparison between schizophrenia and matched controls, which is crucial to tease out treatment- or age-associated effects. A powerful feature of the *Taiji* framework is to allow analysis of individual samples (Zhang et al., 2019). This integrative analysis of transcriptomic and epigenomic data at the systems level uncovered key regulators and important pathways based on their global importance in the genetic networks. We found that transcriptional mechanisms via novel pathways that had not been previously associated with schizophrenia show alterations in AF-schizophrenia subjects, and that these schizophrenia-linked pathways were statistically unaffected in the AT-schizophrenia group, consistent with a potential role of these epigenomic signatures in the clinical efficacy of antipsychotics. These changes appear to impact glutamatergic neurotransmission, IQGAP scaffold, autophagy, and ubiquitin B expression in neurons; and post NMDA receptor activation and long-term potentiation in glial cells. Additionally, our data highlight processes related to key pathways that may represent a consequence of antipsychotic medication, rather than a reversal of the molecular alterations observed in AF-schizophrenia subjects. These pathways suggest the existence of compensatory perturbations that emerge in response to repeated antipsychotic drug administration and ultimately restrain their therapeutic effects (Kurita et al., 2012; Ibi et al., 2017). Among these, alterations in p53 activity were apparent as a consequence of antipsychotic treatment. Based on our ability to individually match schizophrenia and control pairs by age, we also revealed the intriguing observation that the effect of age on TF regulation of gene expression was significantly more pronounced in AT-schizophrenia subjects as compared to AF-schizophrenia subjects and controls.

Related to the effect of antipsychotic treatment, frontal cortex samples of schizophrenia subjects were divided into AF and AT based on postmortem toxicological analysis in both blood and when possible brain samples, which provides information about a longer retrospective drug-free period due to the high liposolubility of antipsychotic medications (Voicu and Radulescu, 2009). However, we cannot fully exclude the possibility of previous exposure to antipsychotic medications in the AF-schizophrenia group, and hence that the epigenetic alterations observed exclusively in the AF-schizophrenia group are a consequence of a potential period of decompensation, which typically occurs following voluntary treatment discontinuation (Liu-Seifert et al., 2005). It is also worth noting that our findings were established by examining the average characteristics of entire NeuN^+^ and NeuN^-^ fractions. Further studies of individual neuronal and glial cell subtypes may yield additional information on the role of cell-type-subpopulations (Lau et al., 2020; Nagy et al., 2020).

## Conclusion

Our ChIP-seq/RNA-seq study in postmortem brain samples from schizophrenia subjects and controls suggests cell-type specific epigenomic differences in individuals with schizophrenia, as well as cellular alterations in signaling pathways potentially involved in either the elimination of schizophrenia-related epigenomic alterations upon antipsychotic drug treatment or the antipsychotic-dependent modulation of alternative epigenetic pathways previously unaffected in the untreated schizophrenia cohort. Building on our data, future research could test the causal role of specific molecular pathways implicated in schizophrenia pathophysiology, as well as the therapeutic versus compensatory or negative side epigenomic outcomes induced by chronic treatment with antipsychotic medications.

## Materials and methods

### Data and code availability

The raw ChIP-seq and RNA-seq data were deposited in dbGaP under accession number phs002487.v1.p1. The processed data can be accessed via Gene Expression Omnibus (GEO) under accession number GSE174407. The code used for the analysis was deposited on GitHub (https://github.com/changlulab/Zhu_Ainsworth_et_al_2023).

### Post-mortem Human Brain Tissue Samples

Human brains were obtained during autopsies performed at the Basque Institute of Legal Medicine, Bilbao, Spain. The study was developed in compliance with policies of research and ethical review boards for post-mortem brain studies (Basque Institute of Legal Medicine, Spain). Deaths were subjected to retrospective searching for previous medical diagnosis and treatment using examiner’s information and records of hospitals and mental health centers. After searching of antemortem information was fulfilled, 29 subjects (Caucasian) who had met criteria of schizophrenia according to the Diagnostic and Statistical Manual of Mental Disorders (DSM-IV) (American Psychiatric Association, 1994) were selected. A toxicological screening for antipsychotics, other drugs and ethanol was performed on blood samples collected at the time of death and, when possible, postmortem brain samples. The toxicological assays were performed at the National Institute of Toxicology, Madrid, Spain, using a variety of standard procedures including radioimmunoassay, enzymatic immunoassay, high-performance liquid chromatography and gas chromatography-mass spectrometry. Controls (Caucasian) for the present study were chosen among the collected brains on the basis, whenever possible, of the following cumulative criteria: (i) negative medical information on the presence of neuropsychiatric disorders or drug abuse; (ii) appropriate sex, age, postmortem delay (time between death and autopsy), and freezing storage time to match each subject in the schizophrenia group; (iii) sudden and unexpected death (motor vehicle accidents); and (iv) toxicological screening for psychotropic drugs with negative results except for ethanol. Specimens of frontal cortex (Brodmann area 9) were dissected at autopsy (0.5-1 g tissue) on an ice-cooled surface and immediately stored at -80°C until use. The schizophrenia subjects were divided into antipsychotic-free and antipsychotic-treated according to the presence or absence of antipsychotics in blood samples at the time of death. The definitive pairs of antipsychotic-free schizophrenics and respective matched controls are shown in Table S1, and the definitive pairs of atypical antipsychotic-treated schizophrenics and respective matched controls are shown in Table S2. Presence or absence of antipsychotic medications was confirmed by toxicological analysis in postmortem brain samples of a selected group of schizophrenia subjects and controls (Table S19). Pairs of schizophrenia and matched controls were processed simultaneously and under the same experimental conditions. Tissue pH values were within a relatively narrow range (control subjects: 6.7 ± 0.08; schizophrenic subjects: 6.6 ± 0.06). Brain samples were also assayed for RIN (RNA integrity number) values using the Agilent 2100 Bioanalyzer (Applied Biosystems) – control subjects: 7.87 ± 0.21; schizophrenic subjects: 7.61 ± 0.32).

### Nuclei isolation and sorting via FACS

Nuclei isolation from frozen tissues (never fixed) of postmortem human brain samples was conducted using a published protocol (Lake et al., 2016) with some modifications. Frontal cortex samples from schizophrenic individuals and individually matched controls were always processed in the same batch. Briefly, all steps were conducted on ice, and all centrifugation was conducted at 4 °C. One piece of brain tissue (∼300 mg) was placed in 3 ml of ice-cold nuclei extraction buffer (NEB) [0.32 M sucrose, 5 mM CaCl_2_, 3 mM Mg(Ac)_2_, 0.1 mM EDTA, 10 mM Tris-HCl, and 0.1%(v/v) Triton X-100] with freshly added 30 µl of protease inhibitor cocktail (PIC, Sigma-Aldrich), 3 µl of 100 mM phenylmethylsulfonyl fluoride (PMSF, Sigma-Aldrich) in isopropyl alcohol, 3 µl of 1 M dithiothreitol (DTT, Sigma-Aldrich), and 4.5 µl of recombinant ribonuclease (RNase) inhibitor (2313A,Takara Bio). The tissue was homogenized in tissue grinder (D9063, Sigma-Aldrich). The homogenate was filtered with a 40 µm cell strainer (22-363-547, Thermo Fisher Scientific) and collected in a 15-ml centrifuge tube. The cell suspension was centrifuged at 1000 RCF at 4 °C for 10 min. The supernatant was discarded, and the pellet was resuspended in 0.5 ml of ice-cold NEB with freshly added 5 µl of PIC, 0.5 µl of PMSF, 0.5 µl of DTT, and 0.75 µl of RNase inhibitor. 500 µl of the sample was mixed with 750 µl of 50%(w/v) iodixanol (a mixture of 4 ml of OptiPrepTM gradient (Sigma-Aldrich) and 0.8 ml of diluent [150 mM KCl, 30 mM MgCl_2_, and 120 mM Tris-HCl]). The mixture was centrifuged at 10,000 RCF at 4 °C for 20 min. Then, the supernatant was removed and 300 µl of 2%(w/v) normal goat serum (50062Z, Life technologies) in Dulbecco’s PBS (DPBS, Life technologies) was added to resuspended the nuclei pellet. To label and separate NeuN^+^ and NeuN^-^ fractions, 6 µl of 2 ng/ml anti-NeuN antibody conjugated with Alexa 488 (MAB377X, Millipore) in DPBS was added into the nuclei suspension. The suspension was mixed well and incubated at 4 °C for 1 hour on a rotator mixer (Labnet). After incubation, the sample was sorted into NeuN^+^ and NeuN^-^ populations using a BD FACSARIATM Flow Cytometer (BD Biosciences). 400 µl of sorted nuclei suspension (NeuN^+^ or NeuN^-^), containing ∼50000 nuclei (for conducting ChIP-seq and input libraries), was added into 600 µl of ice-cold PBS. 200 µl of 1.8 M sucrose solution, 5 µl of 1 M CaCl_2_, and 3 µl of 1 M Mg(Ac)_2_ were added into the mixture. The solution was mixed well and incubated on ice for 15 min. Then, the sample was centrifuged at 1800 RCF at 4 °C for 15 min. The supernatant was discarded and the pellet was resuspended in 110 µl of PBS with freshly added 1.1 µl of PIC and 1.1 µl of PMSF and stored on ice until use.

### Construction of ChIP-seq libraries

Chromatin fragments were prepared by using micrococcal nuclease (MNase) to digest sorted and concentrated nuclei (NeuN^+^/NeuN^-^) following a published protocol (Zhu et al., 2019). 54 µl of chromatin fragments (from 10,000 nuclei) was used in each ChIP assay for producing 2 replicate libraries. Chromatin immunoprecipitation was carried out using multiplexed MOWChIP assay (Zhu et al., 2019) with anti-H3K4me3 (39159, Active Motif) and anti-H3K27ac (39135, Active Motif) antibody. ChIP-seq libraries were prepared using Accel-NGS 2S Plus DNA Library kit (Swift Biosciences) from the purified immunoprecipitated DNA. The library preparation was conducted without knowledge of the brain sample or the type of histone mark. Minor modification was made to the manufacturer’s procedures as detailed below. In the amplification step, instead of adding 10 µl of low EDTA TE buffer into each reactoin, we added the mixture of 7.5 µl of low EDTA TE buffer and 2.5 µl of 20X Evagreen dye to monitor and quantify PCR amplification. The reaction was stopped when the sample’s fluorescence intensity increased by 3000 relative fluorescence units (RFU). Then, 50 µl of the mixture after PCR amplification was transferred into an Eppendorf tube and mixed with 37.5 µl of SPRI select beads. After 5-min incubation at room temperature, the beads went through a cleanup procedure with 80% ethanol. In the end, the DNA library was eluted from the beads into 7 µl of low EDTA TE buffer.

### Construction of RNA-seq libraries

100 µl of sorted nuclei suspension (NeuN^+^ or NeuN^-^) from brain tissue, containing ∼12,000 nuclei for producing 2 replicate libraries, was used for RNA extraction by using the RNeasy Mini Kit (74104, Qiagen) and RNase-Free DNase Set (79254, Qiagen), following the manufacturer’s instruction. Half of the extracted mRNA (from 6,000 nuclei) in 30-µl volume was concentrated by ethanol precipitation and resuspended in 4.6 µl of RNase-free water. mRNA-seq libraries were prepared using Smart-seq2 (Picelli et al., 2013) and a Nextera XT DNA Library Preparation kit (FC-131-1024, Illumina) following the protocol and the manufacturer’s instructions with minor modification. ∼2 ng of mRNA (in 4.6 µl of water) was mixed with 2 µl of 100 mM oligo-dT primer and 2 µl of 10 mM dNTP mix. After being denatured at 72 °C for 3 min, the mRNA solution was immediately placed on ice. Then, 11.4 µl of reverse transcript mix [1 µl of SuperScript II reverse transcriptase (200 U/ml), 0.5 µl of RNAse inhibitor (40 U/ml), 4 µl of Superscript II first-strand buffer, 1 µl of DTT (100mM), 4 µl of 5 M Betaine, 0.12 µl of 1 M MgCl_2_, 0.2 µl of TSO (100 mM), 0.58 µl of nuclease-free water] was mixed with the mRNA solution and the mixture was incubated at 42 °C for 90 min, followed by 10 cycles of (50 °C for 2 min, 42 °C for 2 min). The reaction was finally inactivated at 70 °C for 15 min. 20 µl of first-strand mixture was then mixed with 25 µl of KAPA HiFi HotStart ReadyMix, 0.5 µl of (100 mM) IS PCR primers, 0.5 µl of Evagreen dye, and 4 µl of nuclease-free water. Generated complementary DNA (cDNA) was amplified by incubated at 98 °C for 1 min, followed by 9-11 cycles of (98 °C 15 s, 67 °C 30 s, 72 °C 6 min). After PCR amplification, 50 µl of PCR mix was purified by using 50 µl of SPRIselect beads. ∼600 pg of purified cDNA was used for Nextera XT library preparation. ChIP-seq and RNA-seq library fragment size were measured by using high sensitivity DNA analysis kit (5067-4626, Agilent) on a TapeStation system (2200, Agilent). After this, 18-22 ChIP-seq and RNA-seq libraries were randomly pooled together. Around 15 and 11 million reads were allocated to each ChIP-seq and RNA-seq library, respectively. The concentration of each library was examined by a KAPA library quantification kit (KK4809, Kapa Biosystems), and then the quantified libraries were pooled at 10 nM. The libraries were sequenced by Illumina HiSeq 4000 with single-end 50-nt read.

### ChIP-seq data processing

Raw ChIP-seq reads and input data, were mapped to human genome (GRCh38) using Bowtie2 (2.2.5). Peaks were called using MACS2 (2.2.7.1) using a q-value cutoff of 0.05 for the narrow marks (H3K4me3 and H3K27ac).

### RNA-seq data processing

The human genome (GRCh38) and comprehensive gene annotation were obtained from GENCODE (v29). Quality control of RNA-seq reads including per sequence GC and adapter content was assessed with FastQC. Reads were mapped with STAR (2.7.0f) with soft-clipping (average of 73.8% (+/- 0.08%) reads uniquely mapped for neurons and 69.0% (+/- 0.99%) reads for glia) and quantified with featureCounts (v2.0.1) using the default parameters.

### Differential analysis for ChIP-seq data

The peaks were called using MACS2 (Zhang et al., 2008). The peaks with q-value<0.05 were taken as input for diffBind R package. We first created cell-type-specific consensus peak sets using Diffbind for neurons and glia separately. Using the function of dba.peakset in diffbind (hg38_blacklist_remove, consensus = DBA_REPLICATE, minOverlap = 2), we detected the “high-confidence” peaks if they were identified in both of the technical replicates of the sample (n=58) in either schizophrenia or control groups and then the “high-confidence” peak sets from each sample of the two groups were combined into a master set of consensus peaks for analysis. The raw read counts were extracted using the function of dba.count (hg38_blacklist_remove, summits = FALSE, peaks = consensus_peaks,filter=1, bScaleControl = TRUE, minCount=1, score=DBA_SCORE_TMM_MINUS_FULL) in diffBind, and the peaks with less than 20 reads in over 50% of the samples were removed before differential analysis. DESeq2 R package was used to perform the differential peaks analysis based on the TMM normalized reads to identify differential peaks between schizophrenia and control cohort (adjusted p-value < 0.05). The p-values were adjusted by performing a standard Bonferroni correction. The following covariates were regressed out: demographic covariates (age at death, sex, PMD and diagnosis) and technical covariates (align rate, unique rate, FRiP, NSC, RSC, the number of identified peaks and PBC) by correlating the top 6 principal components with these covariates. We annotated enhancers (defined as identified H3K27ac peaks that have no overlap regions with promoters) to genes using published Hi-C data on neurons and glia (Hu et al., 2021) when possible and the rest of the enhancers were associated with their nearest genes. We annotated H3K4me3 peaks to genes when they overlapped with the promoter regions.

### ChIP-seq annotation and functional enrichment

GREAT analysis (http://great.standford.edu) was performed on differential peaks using the whole genome as background and default basal extension from 5kb upstream to 1kb downstream of the TSS. Significantly enriched Gene Ontology biological processes were identified using the Panther Classification tools using a hypergeometric test.

### Differential analysis for RNA-seq data

We analyzed the bulk RNA-seq data of 29 schizophrenia subjects and 29 controls. The initial step involved filtering out genes with low read counts (less than 20 reads in over 50% of samples). The analysis then employed a two-step method to estimate the technical and biological noise. The first step was identifying the top 10 principal components (PCs) of the dataset. Subsequently, the correlation between each PC and various experimental (alignment rate, unique rate, exon percentage, number of unique mapped reads) and demographic (sex, age at death, PMD, antemortem diagnosis) factors was calculated. Covariates with high correlation to the PCs were included in the analysis to minimize their impact. The analysis was conducted using the ‘DESeq2’ software package, and genes with a false discovery rate (FDR) below 0.05 were identified as differentially expressed.

### ChIP-qPCR assays

After nuclei extraction, MNase digestion and MOWChIP assays (see above), ChIP DNA was eluted to 10 µl of low EDTA TE buffer. 1 µl of ChIP DNA solution was used for qPCR assays with each primer set. The following qPCR primer pairs were used:

Figure S8A: AGG GAC CTG GAA CAT CTT TG (F); CAT CAT CCT CAG AAG GAG TCT G (R)

Figure S8B: TGG AGA TAG GTG GAT GTT AAG C (F); CCA TAT TGA CCC TGG GCT ATT (R)

Figure S8C: ATG CCA ATT AGG CTA TAG ATG CT (F); CTT AAC AGG GCA CTC TCA GTA AT (R)

Figure S8D: AAA GAG CAA GCA GGG ACT T (F); GAT GTA ATA ACG TGG GAG AGA GG (R)

Figure S8E: AGG AGT GGA TAC AGG GAG ATT AG (F); TGT GTA TTC TGT GTC TGG CTT T (R)

Figure S8F: ACC AAC GAA TAC CCT GCT TT (F); AAG GCC TGG CAA CCT TAA T (R)

The following common negative primer set was used in all samples, against which the enrichment of each positive set was calculated:

GCA GAA CCT AGT TCC TCC TTC AAC (F); AGT CAT CCC TTC CTA CAG ACT GAG A (R) qPCR primer sets were ordered from IDT, made to lab ready formulation (100 µM in low EDTA TE buffer). Ready to use stocks of primer sets were made by combining 10 µl each of both forward and reverse primers of the same set with 80 µl of low EDTA TE buffer. 10 µl of iQ SYBR Green Supermix, 1.6 µl of primer stock, 1 µl of ChIP DNA and 7.4 µl of ultrapure water were added to each qPCR well. Reaction was conducted on a CFX96 real-time PCR machine (Bio-Rad) with C1000 thermal cycler base. All PCR assays were performed using the following thermal cycling profile: 95 °C for 10 min followed by 40 cycles of (95 °C for 15 s, 58 °C for 40 s, 72 °C for 30s). Relative fold enrichment of each positive primer (P) against the common negative primer (N) was calculated using the following equation: Enrichment = 2^Cq(N)-Cq(P)^.

### Taiji pipeline

Active regulatory elements were first identified via the overlap of high confidence peaks from H3K27ac with known gene promoter regions (4kbp upstream and 1kbp downstream of the transcription start sites). The distal H3K27ac peaks were assigned to active promoters using the unsupervised learning method EpiTensor, and assigned as an enhancer-promoter interaction if one locus overlapped with the distal peak and the other locus in the pair overlapped with a known promoter. Putative TF binding motifs were curated from the CIS-BP database (Weirauch et al., 2014). Using FIMO’s algorithm (Grant et al., 2011). TFs were identified as having binding sites within 150-bp regions centered around H3K27ac peak summits. 58 unique NeuN^+^ (29 schizophrenia and 29 control) and 58 unique NeuN^-^ (29 schizophrenia and 29 control) network topologies were thus constructed by forming directed edges between TF and their regulatees, if the TF had a predicted binding site in the gene’s promoter or linked enhancer.

### Personalized PageRank (PPR)

The Personalized PageRank (PPR) algorithm was run to measure the global influence of each node. To initialize the networks, node weights were initialized separately in each cell-type *i*, where a gene’s relative expression level is a z-score transformation of its absolute expression, *z_i_* and the node weight for this gene in cell type *i* is then given by *e^zi^*. Edge weights were determined according to the expression level of the parent node TF and the pooled H3K27ac ChIP-seq peak intensity (strength of the TF-gene association) as previously reported (Zhang et al., 2019). The directionality of the topological edges was reversed and the normalized node weights were then used as the seed vector for the PPR calculation. Post convergence, edge directionality was re-reversed.

### EpiSig analysis

To integrate H3K27ac, H3K4me3 and RNA-seq data from two cell types across the postmortem frontal cortex samples from schizophrenia subjects and controls, EpiSig was employed (Ai et al., 2018). This algorithm detects the significant signals from sequencing data in 5kb bins across the whole genome, and then clusters the regions based on the similar epigenomic profiles across all samples.

### EpiSig differential enrichment analysis

A hypergeometric test was applied to all *EpiSig* clusters to assess the enrichment of differential H3K27ac and H3K4me3 peaks and differentially expressed genes. Clusters with FDR < 0.05 were selected and then ranked according to the number of overlapping peaks for each mark. Peaks were then mapped to genes using GREAT with default settings.

### Age correlation analysis

Raw expression, pairwise expression and pairwise TF PPR age correlations were calculated using the Pearson R correlation. Significance was assessed by calculating p-values for the Pearson R correlations using the t-distribution with n-2 degrees of freedom for the respective cohort.

## Acknowledgements

C.L. dedicates this paper to his elder brother Lv Wei who passed away in 2023 after a 30-year battle with schizophrenia. The authors thank the staff members of the Basque Institute of Legal Medicine for their cooperation in the study.

## Author contributions

B.Z., C.L., and J.G.-M. conceived and designed the experiments. R.A., B.Z., Z.W., W.W., C.L., and J.G.-M. analyzed the data and wrote the manuscript. W.W., C.L., and J.G.-M. supervised the research and obtained funding. B.Z., supervised by C.L., performed epigenomic and transcriptomic assays. R.A., supervised by W.W., conducted data analysis and interpreted findings. Z.W., supervised by W.W., and B.Z., supervised by C.L., helped with data analysis. Z.L., supervised by C.L., performed ChIP-qPCR assays. C.D., supervised by C.L., helped with epigenomic and transcriptomic assays. S.S., supervised by C.L. and J.G.-M., helped with nuclear sorting. L.F.C. and J.J.M. obtained and classified postmortem human brain samples. All authors reviewed and approved the final manuscript.

## Supplementary Figure legends

**Figure S1. Distribution of six quality control metrics for ChIP-seq data on H3K4me3 and H3K27ac in NeuN^+^ and NeuN^-^ nuclei from schizophrenia and control groups, respectively.**

(A) Number of uniquely mapped reads obtained by bowtie2 and bedtools.

(B) Fraction of reads in peaks (FRiP) obtained by *featureCounts*.

(C) PCR Bottleneck Coefficient (PBC) obtained by *encodeChIPqc*.

(D) Normalized Strand Coefficient (NSC) obtained by *Phantompeakqualtools*.

(E) Relative Strand Coefficient (RSC) obtained by *Phantompeakqualtools*.

(F) Number of identified peaks by MACS2.

**Figure S2.** Saturation curves on the relationship between the number of sequencing reads and the number of identified peaks with our ChIP-seq data on H3K4me3 and H3K27ac.

**Figure S3. Distribution of four quality control metrics for RNA-seq data in NeuN^+^ and NeuN^-^ nuclei from schizophrenia and control groups.**

(A) Number of uniquely mapped reads.

(B) Aligned rate.

(C) Percentage of reads overlapping exons.

(D) Deduplicate rate (percentage of unique reads)

**Figure S4.** Representative results from FACS sorting demonstrating the separation of neuronal (NeuN^+^) and non-neuronal (NeuN^-^) nuclei using fluorescence-labeled anti-NeuN antibody in postmortem human frontal cortex samples.

**Figure S5.** Comparative bar plots with the expression of neuronal (A and B) and glial (C and D) cell markers in both NeuN^+^ and NeuN^-^ frontal cortex samples from 29 control subjects (A and C) and 29 individuals with schizophrenia (B and D).

**Figure S6. Venn diagrams of the overlap between the identified peaks from our ChIP-seq study (green) and previous datasets (Girdhar et al., 2022) (cyan).**

(A) H3K27ac peak overlap (Mb) between our study (NeuN^+^) and previous datasets (NeuN^+^).

(B) H3K27ac peak overlap (Mb) between our study (NeuN^+^) and previous datasets (bulk).

(C) H3K27me3 peak overlap (Mb) between our study (NeuN^+^) and previous datasets (NeuN^+^).

**Figure S7. Visualization of peak-wise or gene-wise means and variances of ChIP-seq and RNA-seq data, respectively, by voom plots.** The LOWESS curves (red lines) indicate the smooth mean-variance trend.

(A) ChIP-seq data on H3K4me3 from NeuN^+^ nuclei.

(B) ChIP-seq data on H3K4me3 from NeuN^-^ nuclei.

(C) ChIP-seq data on H3K27ac from NeuN^+^ nuclei.

(D) ChIP-seq data on H3K27ac from NeuN^-^ nuclei.

(E) RNA-seq data from NeuN^+^ nuclei.

(F) RNA-seq data from NeuN^-^ nuclei.

**Figure S8. qPCR validation of selected differential ChIP-seq peaks in NeuN^+^ fraction for H3K27ac (A-C) and H3K4me3 (D-F).**

ChIP DNA yielded by MOWChIP assays was examined by qPCR. Primers were designed for each peak (see *Methods*) and amplicon regions were marked by a black bar under genome browser tracks. A common negative locus at gene *AFM* was used for all assays. Relative enrichment of peaks against the negative locus was calculated and shown for all samples. We conducted one ChIP-qPCR assay and two ChIP-seq replicates on each sample.

**Figure S9. Q-Q plots of the corrected p-value from differential peaks at enhancer/promoter regions and differentially expressed genes.**

(A) Differential enhancer peaks from NeuN^+^ nuclei

(B) Differential enhancer peaks from NeuN^-^ nuclei

(C) Differential promoter peaks from NeuN^+^ nuclei

(D) Differential promoter peaks from NeuN^-^ nuclei

(E) Differentially expressed genes from NeuN^+^ nuclei

(F) Differentially expressed genes from NeuN^-^ nuclei

**Figure S10. Enrichment of various GWAS traits in differential enhancers and promoters.** Significance is presented as -log(p values) using LD score regression (ldsc v1.0.1) using genetic loci previously associated with schizophrenia risk (Trubetskoy et al., 2022) as well as other brain and non-brain related traits (Finucane et al., 2018). High values are in red and low values are in blue.

**Figure S11.** UMAP visualization of the feature matrix of enhancers, promoters, and RNA among AF-schizophrenia, AT-schizophrenia and control subjects.

**Figure S12. Differential peak enrichment analysis for antipsychotic-free NeuN+ nuclei in *EpiSig* clusters.**

(A) Percentage of total AF-specific differential H3K27ac peaks in the 22 significantly enriched (FDR<0.05) *EpiSig* clusters. Top 3 clusters 630, 732 (Section 1) and 449 (Section 2) highlighted in red. Differential peaks (all cluster regions for clusters with less than 10 differential peaks) mapped to 166 genes using GREAT with default settings. Histograms of ln[distance to tss] and peak width for 36 H3K27ac differential peaks and 72 regions in cluster 449 (median distance to tss 210 kbp).

(B) Percentage of total AF-specific differential H3K4me3 peaks in the 22 significantly enriched (FDR<0.05) *EpiSig* clusters. Top 3 clusters 582 (Section 6), 270 and 259 (Section 2) highlighted in red. Differential peaks (all cluster regions for clusters with less than 10 differential peaks) mapped to 128 genes using GREAT with default settings. Histograms of ln[distance to tss] and peak width for 19 H3K27ac differential peaks (median distance to tss 186 kbp) and 16 H3K4me3 differential peaks (median distance to tss 141 kbp).

(C) Functional enrichment analysis for the union of gene sets (294 genes) (Table S13).

**Figure S13. Differential peak enrichment analysis for antipsychotic-treated NeuN^+^ nuclei in *EpiSig* clusters.**

(A) Percentage of total AT-specific differential H3K27ac peaks in the 99 significantly enriched (FDR<0.05) *EpiSig* clusters. Top 2 clusters 6 and 8 (Section 4) highlighted in red. Differential peaks mapped to 794 genes using GREAT with default settings. Histograms of ln[distance to tss] and peak width for 623 H3K27ac differential peaks showing bimodal proximal/distal distribution.

(B) Percentage of total AT-specific differential H3K4me3 peaks in the 99 significantly enriched (FDR<0.05) *EpiSig* clusters. Top 2 clusters 79 (Section 3) and 112 (Section 6) highlighted in red. Differential peaks mapped to 80 genes using GREAT with default settings. Histograms of ln[distance to tss] and peak width for 19 H3K27ac differential peaks (median distance to tss 83 kbp) and 37 H3K4me3 differential peaks (median distance to tss 142 kbp).

(C) Functional enrichment analysis for the H3K4me3 differential peak gene set (80 genes) (Table S16).

**Figure S14. Differential peak enrichment analysis for antipsychotic-treated NeuN^-^ nuclei in *EpiSig* clusters.**

(A) Percentage of total AT-specific differential H3K27ac peaks in the 72 significantly enriched (FDR<0.05) *EpiSig* clusters. Top 2 clusters 582 (Section 6) and 703 (Section 1) highlighted in red. Differential peaks mapped to 100 genes using GREAT with default settings. Histograms of ln[distance to tss] and peak width for 61 H3K27ac differential peaks showing bimodal proximal/distal distribution.

(B) Functional enrichment analysis for the H3K27ac differential peak gene set (100 genes) (Table S17).

## Supplementary Tables

**Table S1.** Demographic information of AF-schizophrenia subjects and controls

**Table S2.** Demographic information of AT-schizophrenia subjects and controls

**Table S3.** Quality control measurements on ChIP-seq datasets

**Table S4.** Quality control measurements on RNA-seq datasets

**Table S5.** Transcription frequency in transcripts per kilobase million (TPM) on key marker genes for neuronal and glial nuclei samples from control and schizophrenia subjects

**Table S6.** Differential histone modification peaks and DEGs obtained by comparing schizophrenia and controls

**Table S7.** Overlap of DEGs identified in our study and a previous snRNA-seq study

**Table S8.** Genes identified via differential analyses of enhancers, promoters and RNA by comparing schizophrenia and controls

**Table S9.** *EpiSig* analysis in NeuN^+^ nuclei from the frontal cortex of schizophrenia subjects and controls

**Table S10.** *EpiSig* analysis in NeuN^-^ nuclei from the frontal cortex of schizophrenia subjects and controls

**Table S11.** *Taiji* analysis in NeuN^+^ and NeuN^-^ nuclei from the frontal cortex of schizophrenia subjects and controls

**Table S12.** *Taiji* analysis in NeuN^+^ and NeuN^-^ nuclei from the frontal cortex of AF-schizophrenia subjects, AT-schizophrenia subjects, and controls

**Table S13.** *EpiSig* analysis in NeuN^+^ nuclei from the frontal cortex of AF-schizophrenia subjects and controls

**Table S14.** Differential H3K27ac/H3K4me3 peaks and DEGs obtained by comparing AF-schizophrenia and AT-schizophrenia with their respective matched controls

**Table S15.** Genes altered in AF-but not AT-schizophrenics, and those altered in AT-but not AF-schizophrenics. The lists of genes were obtained by differential analyses of enhancers, promoters, and RNA in AF-schizophrenia, AT-schizophrenia, and their respective controls

**Table S16.** *EpiSig* analysis in NeuN^+^ nuclei from the frontal cortex of AT-schizophrenia subjects and controls

**Table S17.** *EpiSig* analysis in NeuN^-^ nuclei from the frontal cortex of AT-schizophrenia subjects and controls

**Table S18.** Effect of age on chromatin organization in the frontal cortex of AF-schizophrenia subjects, AT-schizophrenia subjects, and controls

**Table S19.** Toxicological analysis in postmortem human brain samples.

