## Supplementary Figs 1-14 for "Antipsychotic-induced epigenomic reorganization in frontal cortex of individuals with schizophrenia"

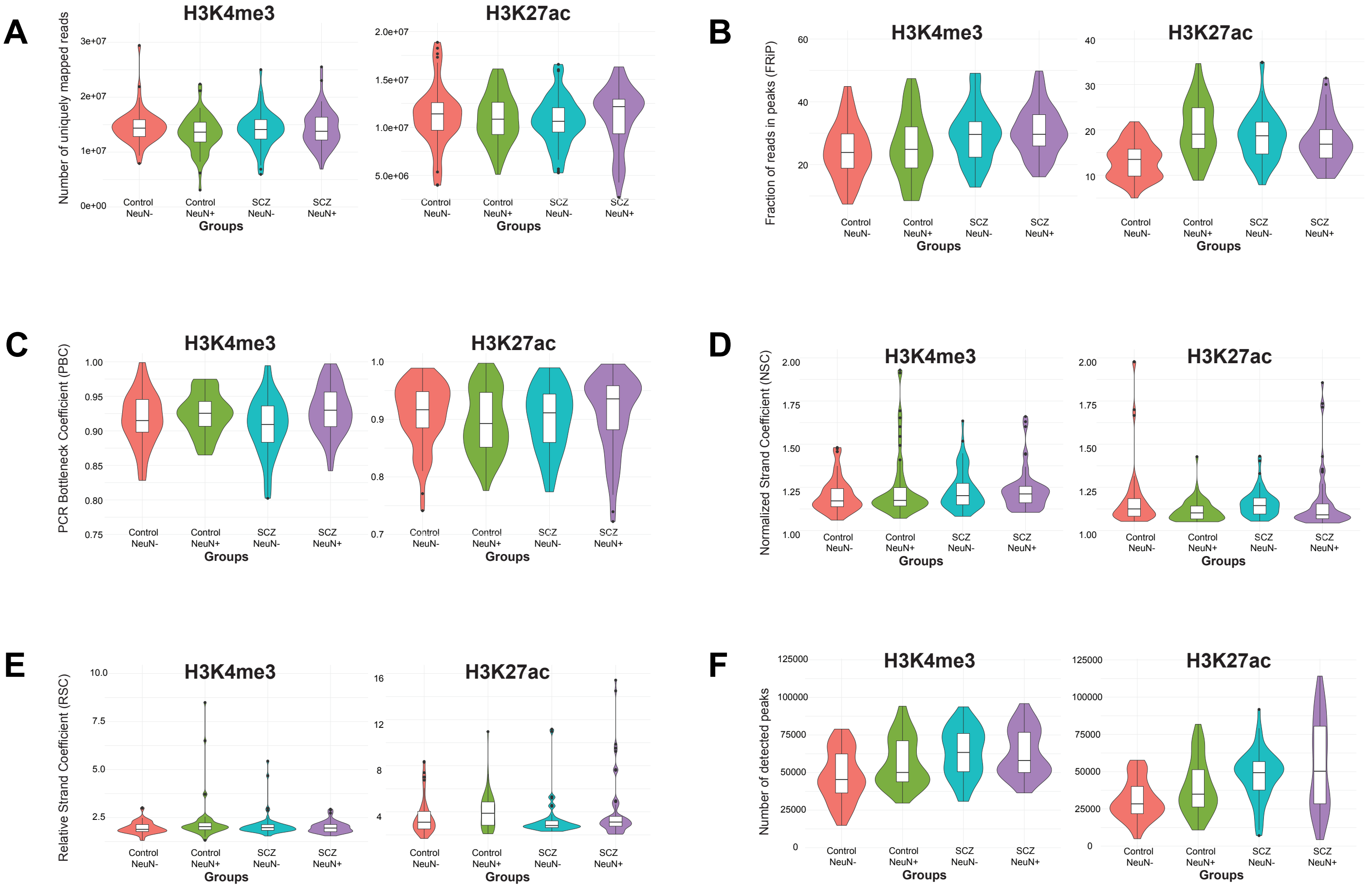


**Zhu, Ainsworth et al (Fig. S1)**

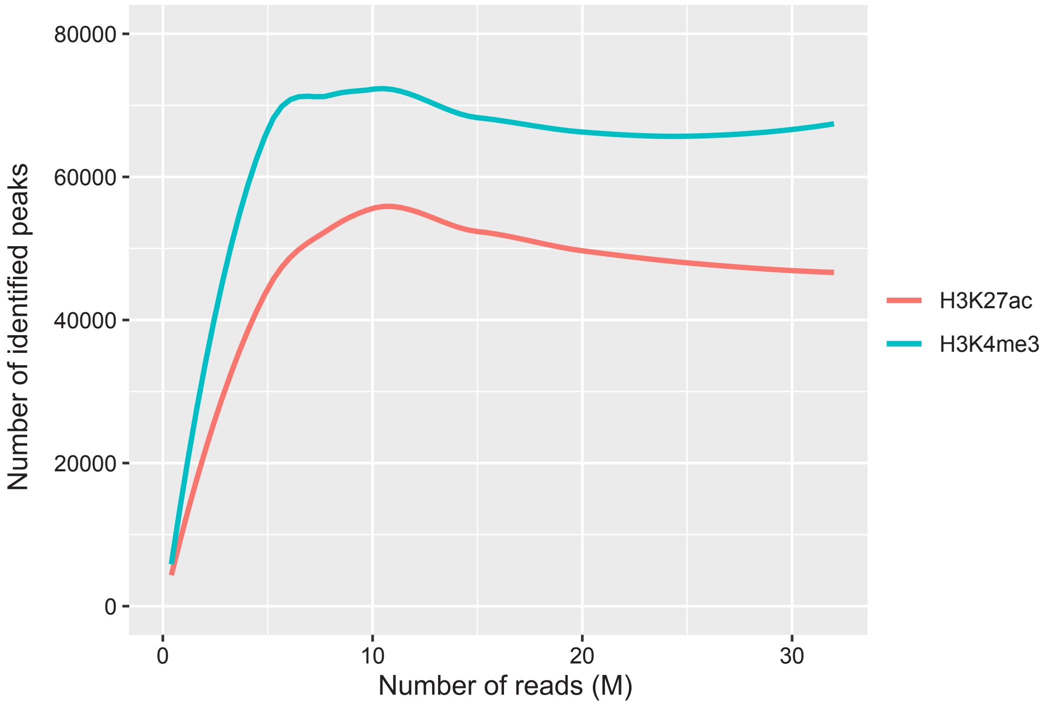


**Zhu, Ainsworth et al (Fig. S2)**

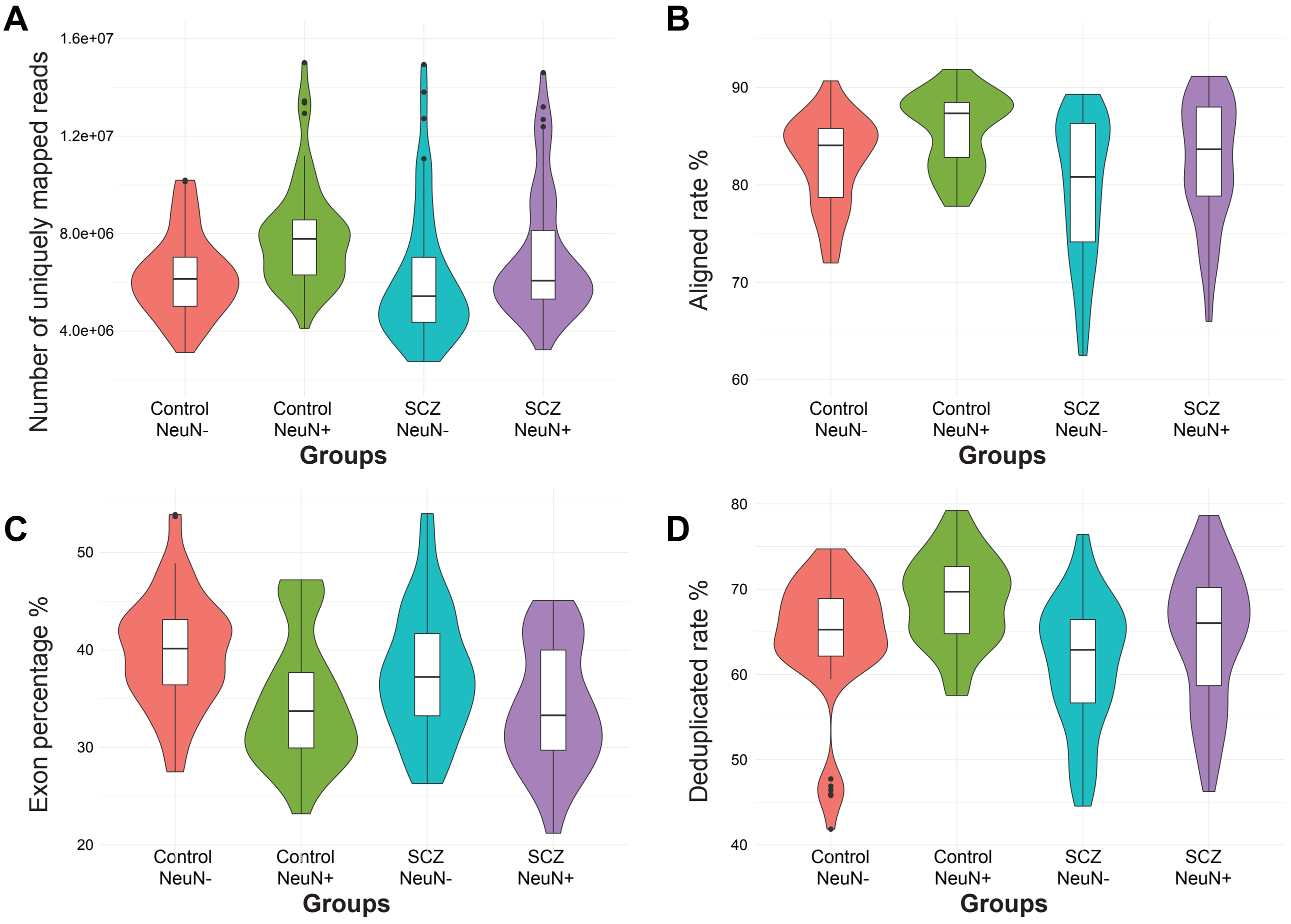


Zhu, Ainsworth et al (Fig. S3)

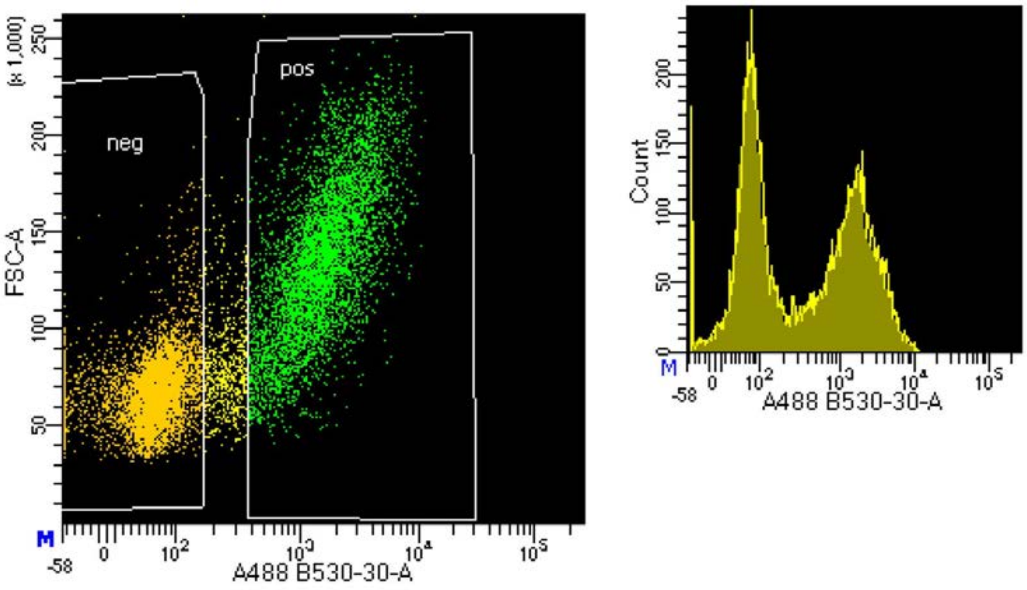


**Zhu, Ainsworth et al (Fig. S4)**

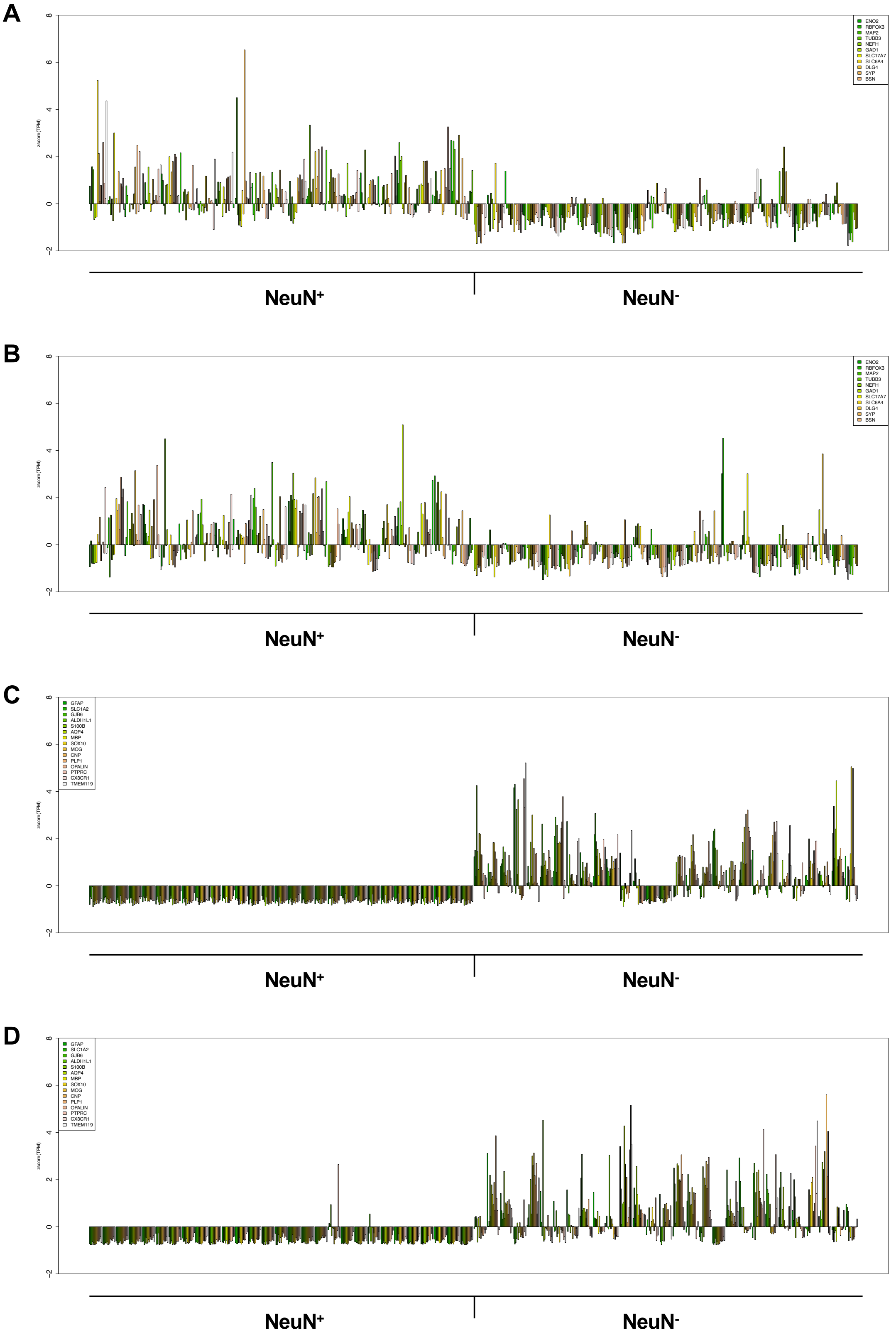


**Zhu, Ainsworth et al (Fig. S5)**

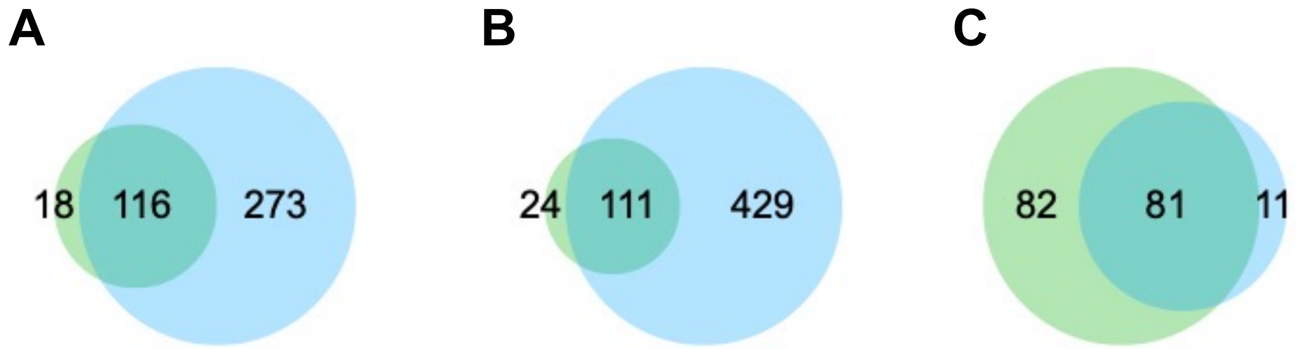


Zhu, Ainsworth et al (Fig. S6)

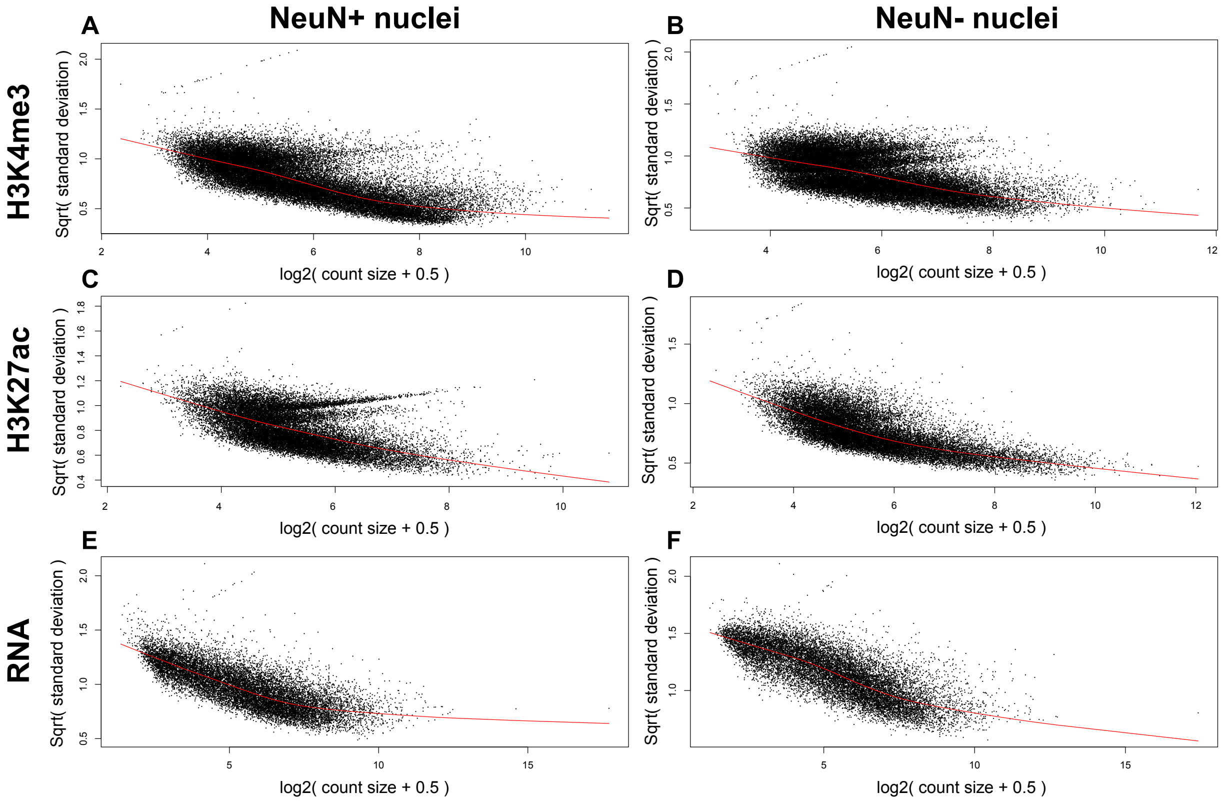


Zhu, Ainsworth et al (Fig. S7)

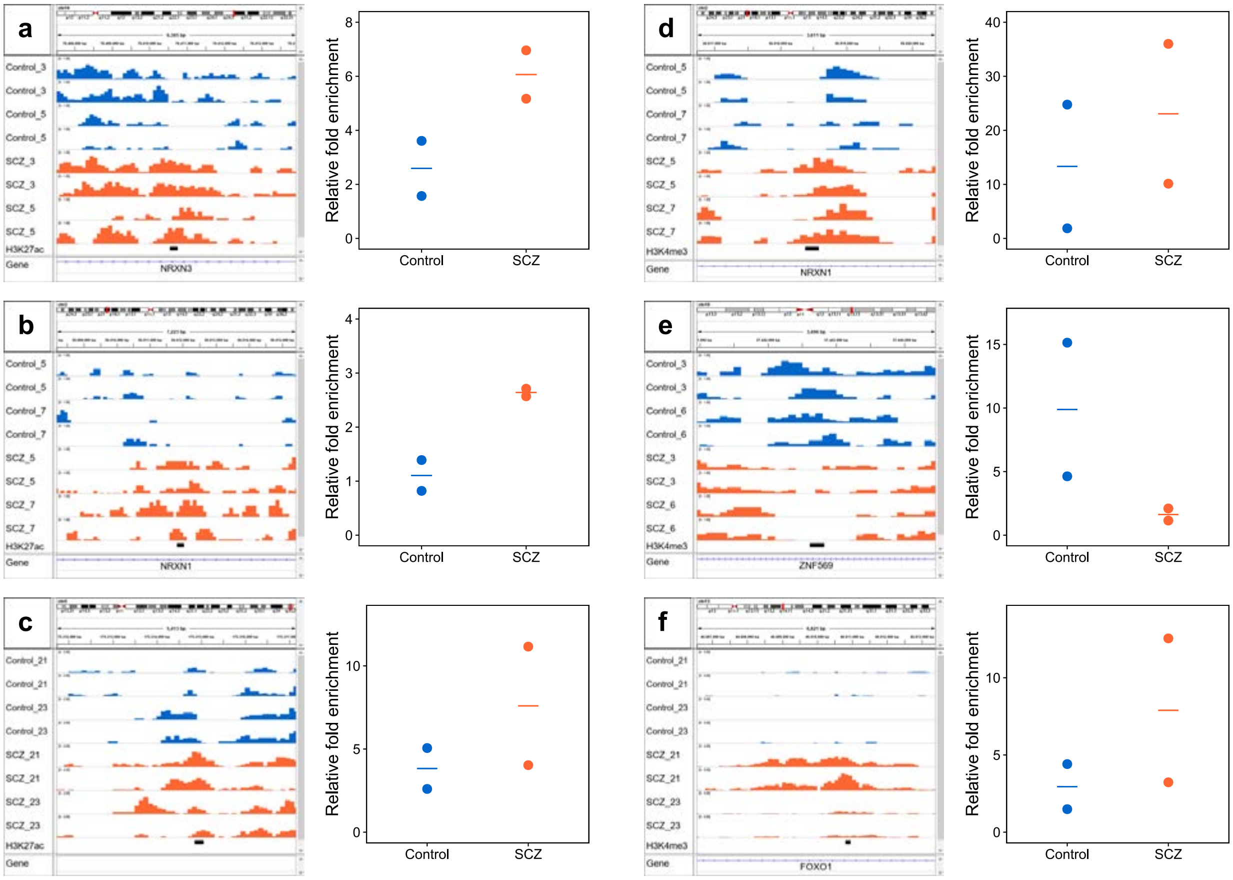


**Zhu, Ainsworth et al (Fig. S8)**

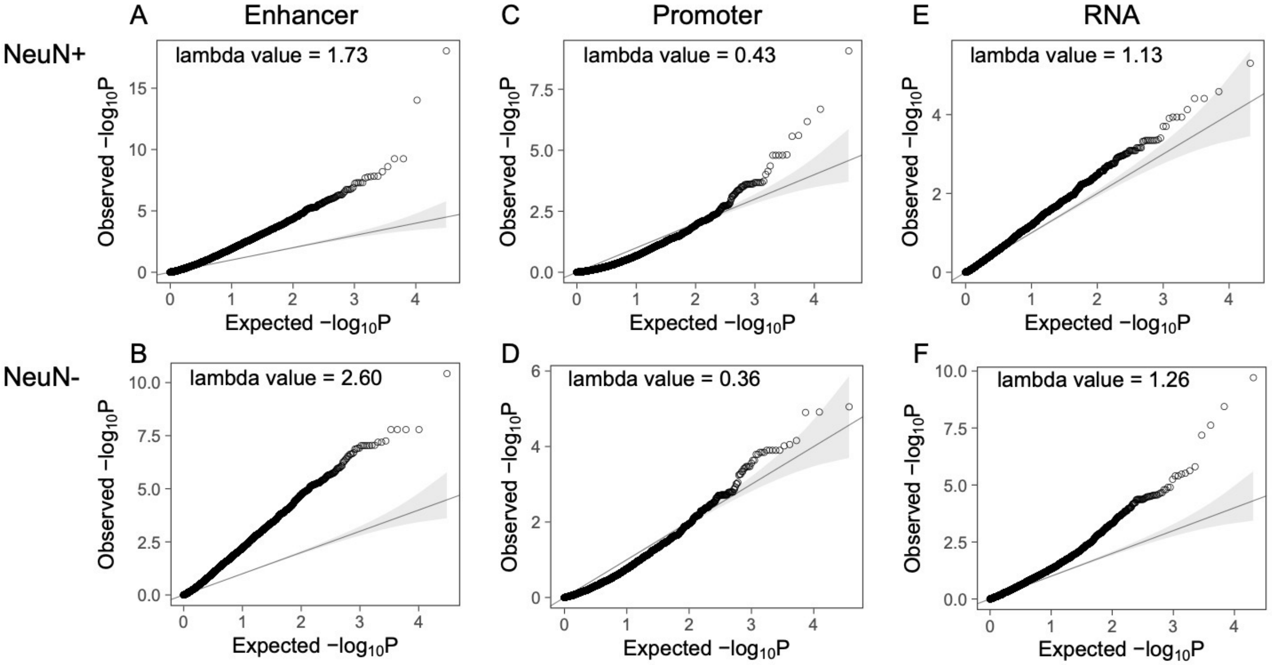


Zhu, Ainsworth et al (Fig. S9)

| | Diff. enhancer NeuN+ | Diff. enhancer NeuN- | Diff. promoter NeuN+ | Diff. promoter NeuN- |
|---|---|---|---|---|
| **Schizophrenia** | 22.47 | 10.12 | 2.49 | 3.75 |
| **Depression** | 5.88 | 5.24 | 1.08 | 1.62 |
| **Neuroticism** | 4.16 | 4.27 | 0.04 | 1.39 |
| **ADHD** | 3.63 | 1.93 | 0.35 | 0.56 |
| **Alzheimer** | 3.35 | 0.55 | 0.15 | 1.34 |
| **Bipolar disorder** | 2.09 | 1.71 | 0.60 | 0.68 |
| **Coronary artery disease** | 1.74 | 1.37 | 0.03 | 1.00 |
| **Autism** | 1.69 | 0.07 | 0.23 | 0.34 |
| **Ulcerative Colitis** | 1.49 | 0.53 | 0.45 | 2.19 |
| **Crohns Disease** | 1.40 | 2.86 | 0.40 | 1.89 |
| **Epilepsy** | 1.12 | 0.38 | 0.32 | 0.58 |
| **Type 2 Diabetes** | 0.98 | 2.04 | 0.06 | 0.12 |
| **Celiac** | 0.53 | 1.99 | 0.59 | 0.99 |

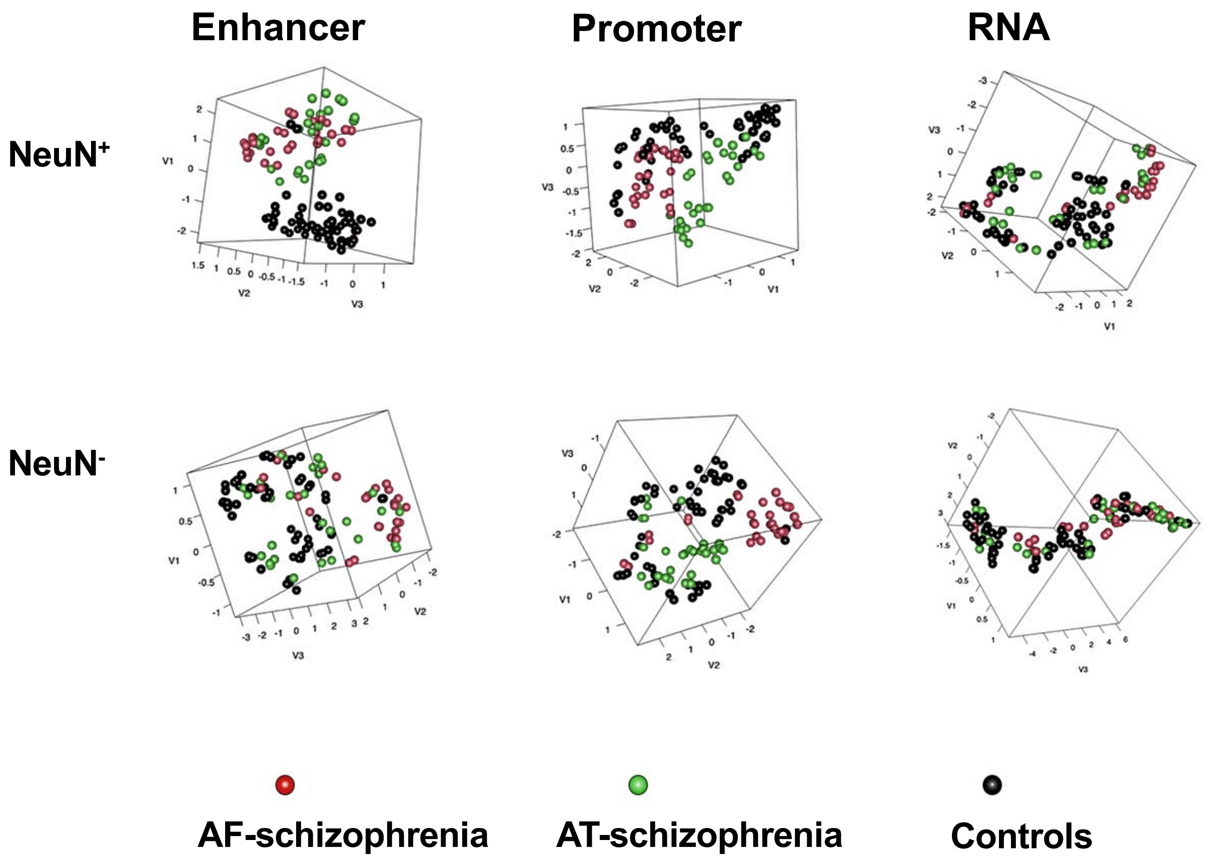


Zhu, Ainsworth et al (Fig. S11)

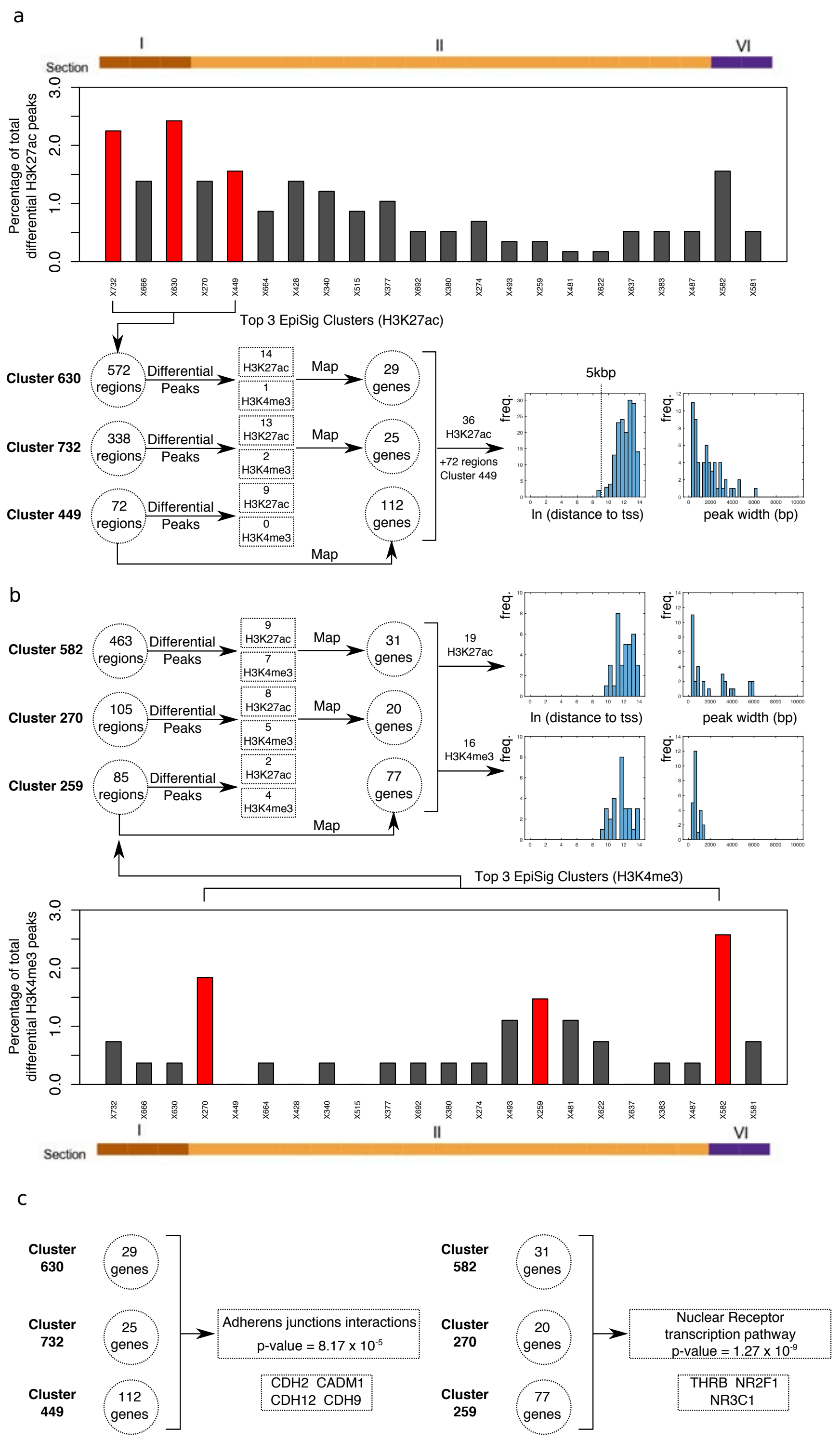


Zhu, Ainsworth et al (Fig. S12)

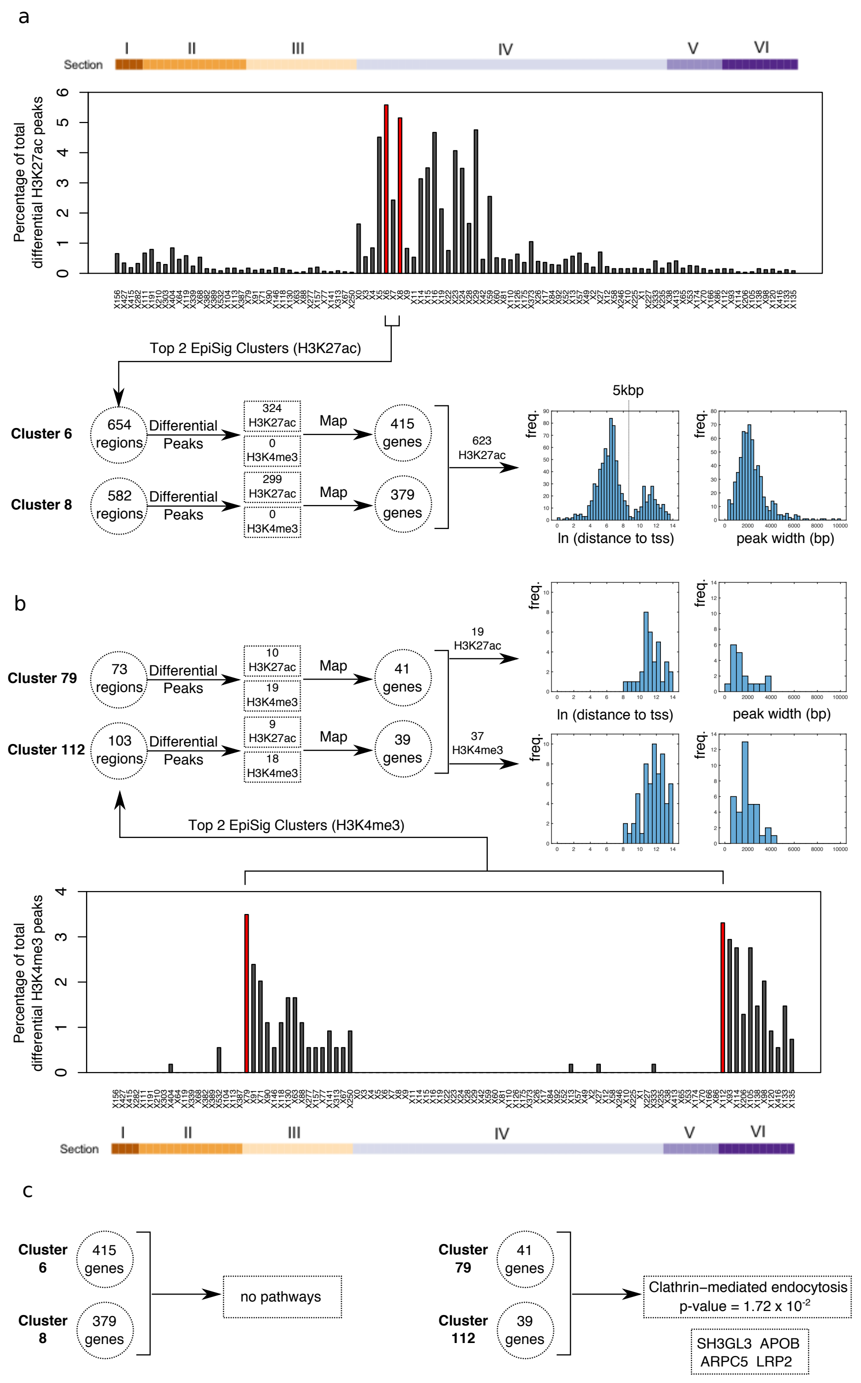


**Zhu, Ainsworth et al (Fig. S13)**

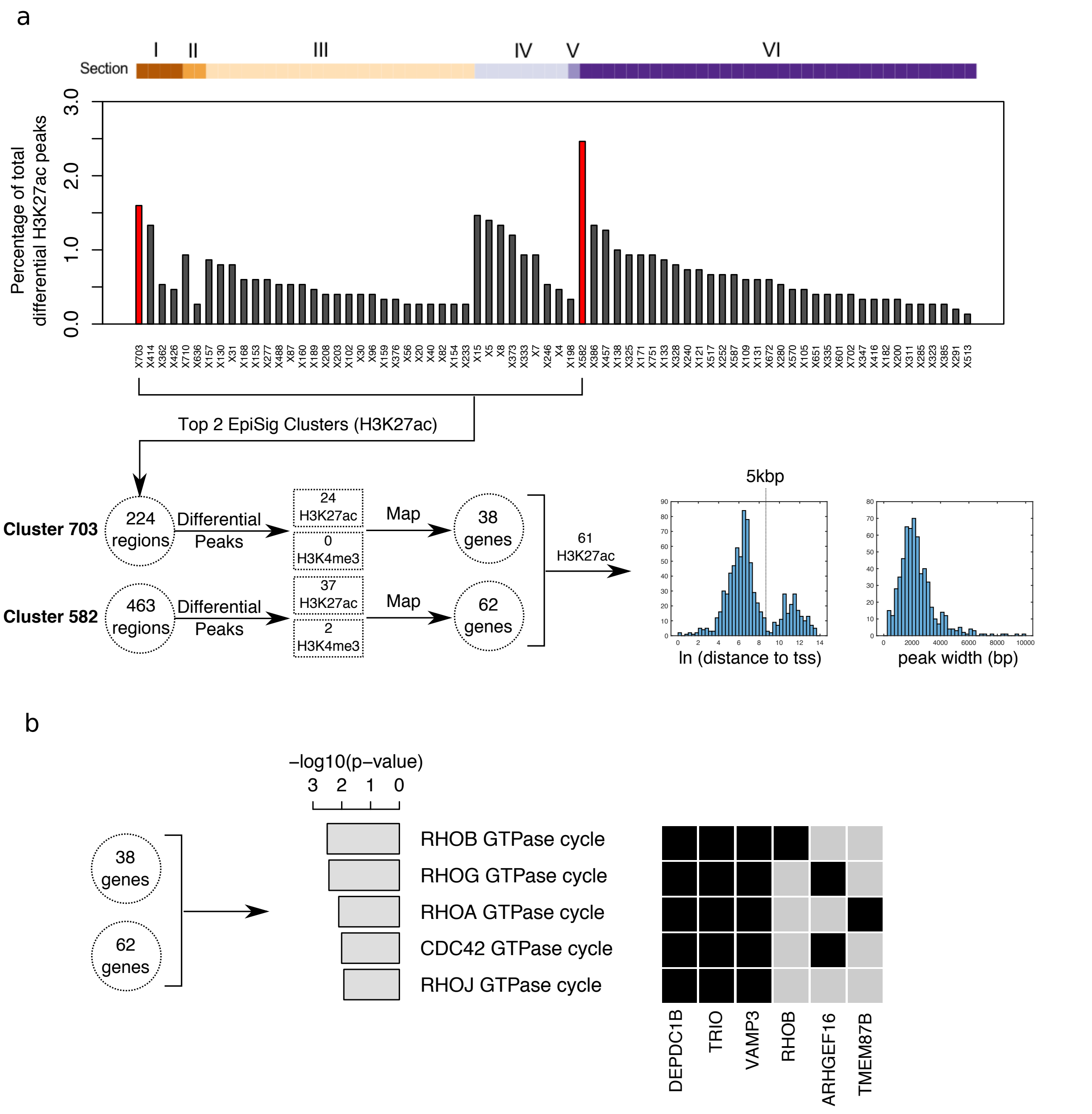


Zhu, Ainsworth et al (Fig. S14)
