## Supplementary material for "Antipsychotic-induced epigenomic reorganization in frontal cortex of individuals with schizophrenia": TableS1

**Table S1.** Demographic characteristics of antipsychotic-free schizophrenia subjects and their respective control subjects.

|  | ***Sex***  ***(F/M)*** | ***Age at death***  ***(years)*** | ***Postmortem delay (h)*** | Antipsychotics in blood | ***Additional drugs in blood*** |
| --- | --- | --- | --- | --- | --- |
| Control 1 | M | 22 | 20 |  | (-) |
| Schizophrenia 1 | M | 24 | 45 | (-) | (-) |
| Control 2 | M | 21 | 30 |  | ETH (0.34 g/l) |
| Schizophrenia 2 | M | 21 | 24 | (-) | (-) |
| Control 3 | M | 18 | 25 |  | (-) |
| Schizophrenia 3 | M | 18 | 18 | (-) | (-) |
| Control 4 | M | 32 | 28 |  | ETH (0.68 g/l) |
| Schizophrenia 4 | M | 31 | 14 | (-) |  |
| Control 5 | F | 28 | 55 |  | (-) |
| Schizophrenia 5 | F | 28 | 22 | (-) | (-) |
| Control 6 | M | 28 | 30 |  | (-) |
| Schizophrenia 6 | M | 27 | 24 | (-) | (-) |
| Control 7 | M | 25 | 20 |  | cannabis |
| Schizophrenia 7 | M | 25 | 16 | (-) | (-) |
| Control 8 | F | 66 | 17 |  | (-) |
| Schizophrenia 8 | F | 67 | 22 | (-) | (-) |
| Control 9 | M | 34 | 17 |  | (-) |
| Schizophrenia 9 | M | 34 | 23 | (-) | (-) |
| Control 10 | F | 51 | 10 |  | (-) |
| Schizophrenia 10 | F | 53 | 18 | (-) | (-) |
| Control 11 | M | 33 | 23 |  | (-) |
| Schizophrenia 11 | M | 32 | 21 | (-) | (-) |
| Control 12 | M | 44 | 23 |  | (-) |
| Schizophrenia 12 | M | 43 | 23 | (-) | (-) |
| Control 13 | M | 49 | 20 |  | (-) |
| Schizophrenia 13 | M | 49 | 19 | (-) | (-) |
| Control 14 | M | 71 | 9 |  | (-) |
| Schizophrenia 14 | M | 70 | 22 | (-) | BDZ |
| Control 15 | F | 74 | 20 |  | (-) |
| Schizophrenia 15 | F | 74 | 30 | (-) | (-) |
| Control | 4F/11M | 39.7 ± 4.8 | 23.1 ± 2.7 |  |  |
| Schizophrenia | 4F/11M | 39.7 ± 4.8 | 22.7 ± 1.8 |  |  |

Antipsychotics were not detected in blood samples of schizophrenia subjects. All schizophrenia subjects included, except schizophrenia 8, schizophrenia 10, schizophrenia 13, and schizophrenia 15, committed suicide. Abbreviations: benzodiazepines (BDZ). Ethanol in blood is coded as ETH.
