## Supplementary material for "Antipsychotic-induced epigenomic reorganization in frontal cortex of individuals with schizophrenia": TableS2

**Table S2.** Demographic characteristics of antipsychotic-treated schizophrenia subjects and their respective control subjects.

|  | ***Sex***  ***(F/M)*** | ***Age at death***  ***(years)*** | ***Postmortem delay (h)*** | Antipsychotics in blood | ***Additional drugs in blood*** |
| --- | --- | --- | --- | --- | --- |
| Control 16 | M | 32 | 20 |  | ETH (2.37 g/l) |
| Schizophrenia 16 | M | 32 | 8 | QTP | BDZ |
| Control 17 | M | 26 | 27 |  | (-) |
| Schizophrenia 17 | M | 26 | 39 | OLZ | (-) |
| Control 18 | M | 36 | 18 |  | ETH (1.69 g/l) |
| Schizophrenia 18 | M | 35 | 11 | CLZ | BDZ |
| Control 19 | M | 52 | 23 |  | (-) |
| Schizophrenia 19 | M | 51 | 28 | OLZ, CLT | (-) |
| Control 20 | M | 56 | 24 |  | (-) |
| Schizophrenia 20 | M | 58 | 24 | PAL | (-) |
| Control 21 | F | 48 | 21 |  | (-) |
| Schizophrenia 21 | F | 52 | 31 | RIS | (-) |
| Control 22 | M | 34 | 36 |  | ETH (0.74 g/l) |
| Schizophrenia 22 | M | 34 | 21 | HAL | BDZ |
| Control 23 | M | 42 | 6 |  | BDZ (0.72 g/l) |
| Schizophrenia 23 | M | 43 | 6 | PAL | BDZ |
| Control 24 | F | 50 | 10 |  | (-) |
| Schizophrenia 24 | F | 50 | 14 | RIS, QTP | BDZ |
| Control 25 | F | 60 | 48 |  | (-) |
| Schizophrenia 25 | F | 60 | 22 | CLZ, ASU | BDZ |
| Control 26 | M | 54 | 16 |  | ETH (0.58 g/l) |
| Schizophrenia 26 | M | 56 | 12 | OLZ, CLT | (-) |
| Control 27 | M | 41 | 14 |  | (-) |
| Schizophrenia 27 | M | 41 | 11 | QTP, CLT | BDZ |
| Control 28 | M | 36 | 12 |  | ETH (1.0 g/l) |
| Schizophrenia 28 | M | 36 | 8 | OLA | (-) |
| Control 29 | F | 41 | 22 |  | (-) |
| Schizophrenia 29 | F | 42 | 14 | CLZ, SUL, QTP | (-) |
| Control | 4F/10M | 43.4 ± 2.7 | 21.2 ± 2.8 |  |  |
| Schizophrenia | 4F/10M | 44.0 ± 2.8 | 17.8 ± 2.6 |  |  |

Therapeutic levels of amisulpiride (ASU), clotiapine (CLT), clozapine (CLZ), haloperidol (HAL), olanzapine (OLZ), quetiapine (QTP), risperidone (RIS), paliperidone (PAL), and sulpiride (SUL) were detected in blood samples of schizophrenia subjects. All schizophrenia subjects included, except schizophrenia 18, schizophrenia 19, schizophrenia 20, schizophrenia 22, schizophrenia 25, and schizophrenia 26, committed suicide. Abbreviations: benzodiazepines (BDZ). Ethanol in blood is coded as ETH.
